# scGPT: Towards Building a Foundation Model for Single-Cell Multi-omics Using Generative AI

**DOI:** 10.1101/2023.04.30.538439

**Authors:** Haotian Cui, Chloe Wang, Hassaan Maan, Kuan Pang, Fengning Luo, Bo Wang

## Abstract

Generative pre-trained models have achieved remarkable success in various domains such as natural language processing and computer vision. Specifically, the combination of large-scale diverse datasets and pre-trained transformers has emerged as a promising approach for developing foundation models. Drawing parallels between linguistic constructs and cellular biology — where texts comprise words, similarly, cells are defined by genes — our study probes the applicability of foundation models to advance cellular biology and genetics research. Utilizing the burgeoning single-cell sequencing data, we have pioneered the construction of a foundation model for single-cell biology, scGPT, which is based on generative pre-trained transformer across a repository of over 33 million cells. Our findings illustrate that scGPT, a generative pre-trained transformer, effectively distills critical biological insights concerning genes and cells. Through the further adaptation of transfer learning, scGPT can be optimized to achieve superior performance across diverse downstream applications. This includes tasks such as cell-type annotation, multi-batch integration, multi-omic integration, genetic perturbation prediction, and gene network inference. The scGPT codebase is publicly available at https://github.com/bowang-lab/scGPT.

## 1 Main

Single-cell RNA sequencing (scRNA-seq), by enabling intricate characterization of distinct cell types and advancing our understanding of disease pathogenesis, paves the way for cellular heterogeneity exploration, lineage tracking, pathogenic mechanism elucidation, and, ultimately, personalized therapeutic strategies [1, 2, 3, 4]. The broad-scale application of scRNA-seq has led to comprehensive data atlases like the Human Cell Atlas, which now encompasses tens of millions of cells [5, 6, 7, 8]. Furthermore, such “omic” data is expanding exponentially. Recent sequencing technology advancements in the diversity of data modalities extend our understanding beyond genetics to epigenetics, transcriptomics, and proteomics, thus providing multi-modal insights [9, 10]. These breakthroughs have also raised new research questions such as reference mapping, perturbation prediction, and multi-omic integration [11, 12, 13, 14, 15]. Given the rapid expansion of sequencing data, it is critical to parallelly develop methodologies capable of effectively harnessing, enhancing, and adapting to these burgeoning developments.

One promising approach to address this challenge is the generative pre-training of foundation models [16, 17]. Generative pre-training has recently achieved unprecedented success across various fields by learning from extensive datasets. The most well-known applications include computer vision and natural language generation (NLG) [18, 19, 20]. These foundation models such as DALL-E2 and GPT-4 follow a paradigm of pre-training transformers on large-scale diverse datasets [19, 20] that can be readily customized for a variety of downstream tasks and scenarios. More interestingly, these generative pre-trained models consistently outperform task-specific models trained from scratch [21, 22, 23]. This indicates a task-agnostic understanding of knowledge in these domains, inspiring us to explore its adoption for sing-cell omics research. However, current machine-learning-based methods in single-cell research are rather scattered, with specific models dedicated to distinct analysis tasks [24, 25, 26]. As a result, the datasets used in each study are often limited in breadth and scale [8]. To confront this limitation, there is a need for a foundation model that is pre-trained on large-scale data and can comprehend the complex interactions between genes across diverse tissues. We expect that such a model would provide a solid foundation and contribute to the discovery of new biological insights by leveraging the knowledge learned from millions of sequenced cells.

To enhance the modeling of large-scale data of single-cell sequencing, we draw inspiration from the self-supervised pre-training workflow in NLG. The self-attention transformer has been verified as an effective and efficient architecture to model input tokens of words [27]. While texts are made up of words, cells can be characterized by genes and the protein products they encode. By learning gene and cell embeddings simultaneously, akin to word and sentence embeddings in NLG, we can better comprehend the characteristics of cells based on the genes they express. Moreover, the flexible nature of transformer input tokens enables easy incorporation of additional features and meta information.

In this work, we present the first attempt to build a single-cell foundation model, scGPT, by generative pre-training on over 33 million cells. We introduce new techniques to address the methodology and engineering challenges of pre-training on large-scale single-cell omic data. To handle large-scale data, we use an in-memory data structure that allows fast access to store hundreds of datasets. We establish a unified generative pre-training workflow specifically for the non-sequential omic data, and adapt the transformer architecture to simultaneously learn cell and gene representations. Additionally, we provide common pipelines with task-specific objectives for model fine-tuning, designed to facilitate the application of the pre-trained model across a range of downstream tasks.

Our model, scGPT, demonstrates the transformative potential of the single-cell foundation model through three key aspects. First, scGPT represents the first large-scale *generative* foundation models that enable transfer learning across a diverse range of downstream tasks. By achieving state-of-the-art performance on cell type annotation, genetic perturbation prediction, batch correction, and multi-omic integration, we showcase the effectiveness of the “pre-training universally, finetuning on demand” approach as a generalist solution for computational applications in single-cell omics. Notably, scGPT is the only foundation model that can integrate multiple singlecell omics including scATAC-seq data. Second, through the comparison of gene embeddings and attention weights between the fine-tuned and raw pre-trained models, scGPT uncovers valuable biological insights into gene-gene interactions specific to various conditions, such as cell types and perturbation states. Third, our observations reveal a scaling law: larger pre-training data sizes yield superior pre-trained embeddings and further lead to improved performance on downstream tasks. This finding highlights the exciting prospect that foundation models can continuously improve alongside the expansion of available sequencing data in the research community. Based on these findings, we envision that the adoption of pre-trained foundation models will greatly expand our understanding of cellular biology and serve as a solid foundation for future discoveries. The release of the scGPT models and workflow aims to empower and expedite research in these areas and beyond.

## 2 Results

### 2.1 Single-cell transformer foundation model overview

Single-cell sequencing enables the profiling of molecular characteristics at the individual cell level. For instance, scRNA-seq measures the abundance of RNA transcripts, providing insights into cell identity, developmental stage, and functionality. We introduce scGPT as the first foundation model in the single-cell domain with a generative pre-training approach. The core model contains stacked transformer layers with multi-head attention [27] that generate cell and gene embeddings simultaneously (Online Methods 4.2). scGPT consists of two stages: the initial general-purpose pre-training on large cell atlases and the follow-up fine-tuning on smaller datasets for specific applications (Figure 1A-C). In the pre-training stage, we introduce a specially designed attention mask and generative training pipeline to train scGPT in a self-supervised manner to jointly optimize cell and gene representations (Online Methods 4.3). This innovative technique successfully addresses the non-sequential nature of gene expression to adapt to the NLG framework of sequential prediction. During training, the model gradually learns to generate gene expression of cells based on cell states or gene expression cues. In the fine-tuning stage, the pre-trained model can be adapted to new datasets and specific tasks (Online Methods 4.5). We offer flexible fine-tuning pipelines suitable for a variety of essential downstream tasks in single-cell research, including scRNA-seq integration with batch correction, cell type annotation, multi-omic integration, perturbation prediction, and gene regulatory network inference. These pipelines enable researchers to leverage the power of scGPT for diverse applications in single-cell analysis.

**Figure 1:**
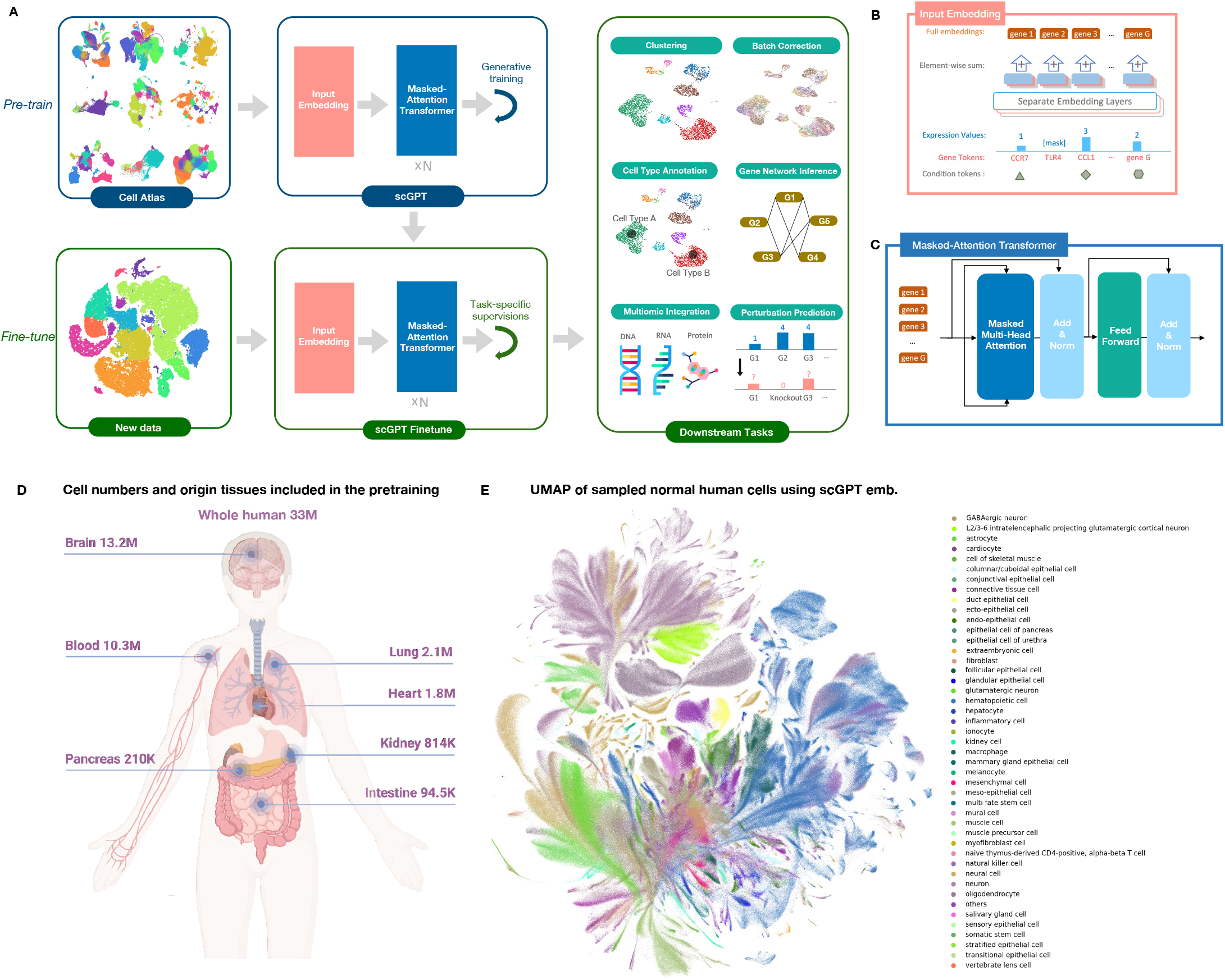
Model Schematic. *(A)* The workflow of scGPT. The model is generatively pre-trained on large-scale scRNA-seq data from cell atlases. For downstream applications, the pre-trained model parameters can be fine-tuned on new data. The core component of scGPT contains stacked transformer blocks with specialized attention masks for generative training. We applied scGPT on a variety of tasks including cell type annotation, batch correction, multi-omic integration, genetic perturbation prediction, and gene network inference. *(B)* The detailed view of the input data embeddings. The input contains three layers of information, gene tokens, expression values, and condition tokens (modality, batch, perturbation conditions, etc.). *(C)* The detailed view of the scGPT transformer layer. We introduced a specially designed attention mask in the Masked Multi-Head Attention block to conduct generative pre-training on single-cell sequencing data. *(D)* The diagram illustrating the size of training data and the organs of origin. The scGPT whole-human model was pre-trained on the scRNA-seq data of 33 million normal human cells. *(E)* UMAP visualization of the pre-trained scGPT cell embeddings (a random 10% subset), colored by major *cell types*.

To collect diverse and extensive sequencing data for the self-supervised pre-training of scGPT, we assembled 33 million scRNA-seq data of human cells under normal (non-disease) conditions, obtained via the CELLxGENE collection [28] (Figure 1D). This comprehensive dataset encompasses a wide range of cell types from 51 organs/tissues and 441 studies, providing a rich representation of cellular heterogeneity across the human body. After pre-training, we visualized the scGPT cell embeddings on 10% of the human cells out of the 33 million data using UMAP visualization [29] (Figure 1E). The resulting UMAP plot exhibits intriguing clarity, with cell types accurately represented by distinct colors at localized regions and clusters. Considering the inclusion of over 400 studies in the dataset, this demonstrates the remarkable capability of pre-training to mitigate technical batch effects.

### 2.2 scGPT improves the precision of cell type annotation

Cell type annotation is a crucial step in single-cell analysis, enabling the identification and characterization of distinct cell populations within sequenced tissues. While several methods have been proposed for cell annotation [30, 31, 32], they often require dimension reduction prior to model input, which can lead to information loss. In contrast, scGPT’s transformer model offers an advantage by directly accepting gene expression as input without the need for prior dimension reduction. This approach provides improved reliability and accuracy as demonstrated in our subsequent analyses.

To evaluate the performance of scGPT for cell-type annotation, we conducted extensive experiments on a variety of diverse datasets. Firstly, we adapted scGPT to predict cell types in a human pancreas (hPancreas) dataset. The model was fine-tuned on a reference data partition and used to predict on a held-out query data partition. We visualized the predictions in Figure 2A. Notably, scGPT achieved high precision (*>* 0.8) for most cell types shown in the confusion matrix (Figure 2B), only except for rare cell types with extremely low cell numbers in the reference partition. For example, fewer than 50 cells belong to the mast and MHC class II cell types out of the 106,000 cells in the reference set. Figure 2C visualizes the cell embeddings in the fine-tuned scGPT, which demonstrate high intra-cell-type similarities.

**Figure 2:**
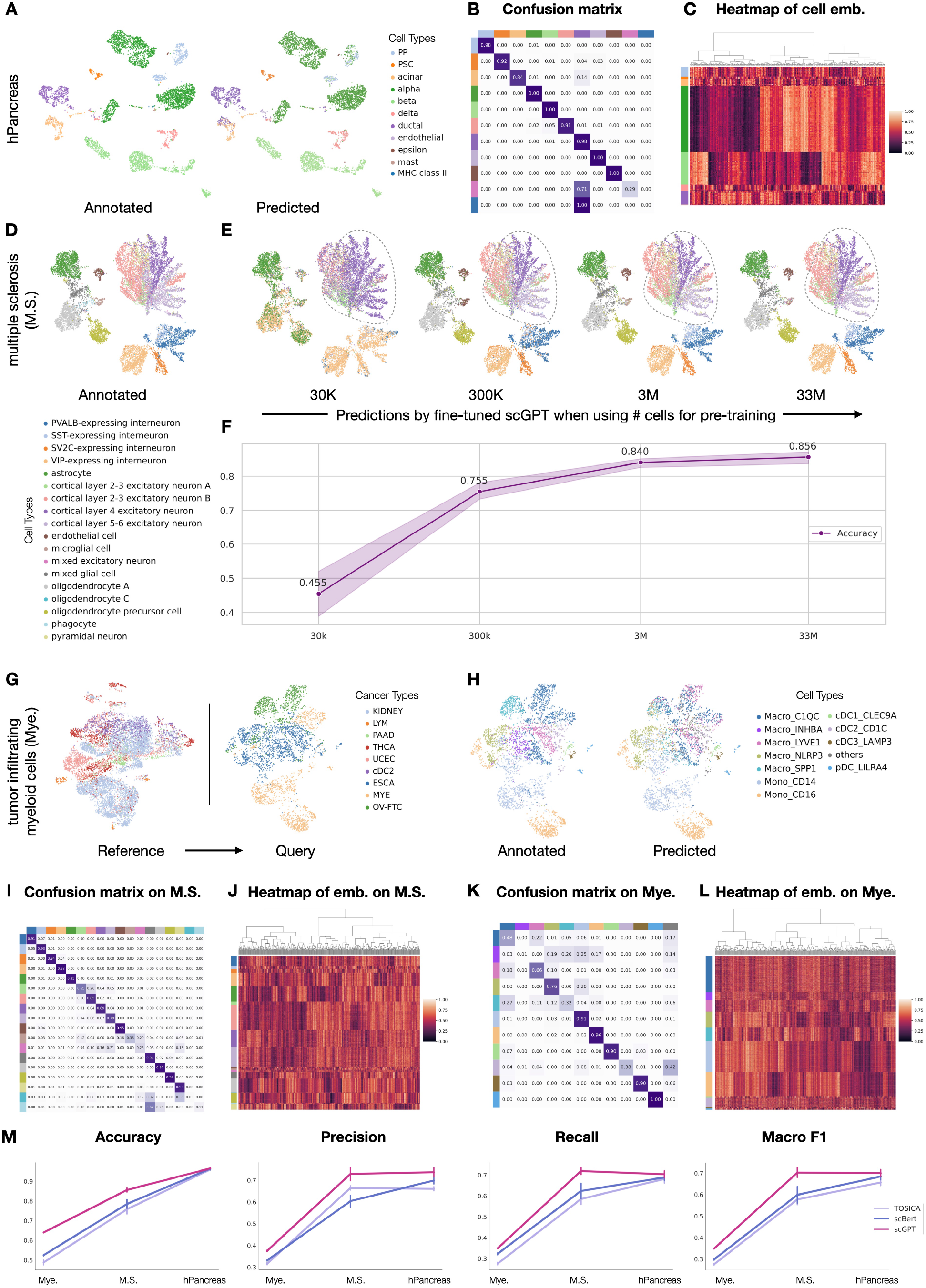
Cell type annotation results using scGPT. *(A)* UMAP of the gene expression of cells from the hPancreas dataset, colored by the *cell types* annotated in the original study (left) and by the *cell types* predicted by the fine-tuned scGPT (right). On the hPancreas dataset, *(B)* The confusion matrix between predicted and annotated cell types, *(C)* the heatmap of the 512-dimensional cell embeddings in scGPT. *(D)* UMAP of the multiple sclerosis (M.S.) dataset colored by the *cell types* annotated in the original study. *(E)* UMAP of M.S. colored by the predictions of four fine-tuned scGPT models using different pre-training models. From left to right, 30K, 300K, 3M, and 33M normal human cells were used during pre-training. *(F)* The prediction accuracy increases as the number of cells grows in pre-training. *(G)* UMAP visualization of the tumor-infiltrating myeloid dataset (Mye.), colored by *cancer types*. scGPT was fine-tuned on the reference partition (left) and evaluated on the query partition (right). These two data partitions contain distinct cancer types. *(H)* UMAP colored by the cell types annotated in the original study (left) and by the scGPT predicted *cell types. (I,K)* The confusion matrices between predicted cell types and actual annotations on M.S. and Mye., respectively. *(J,L)* Heatmaps showing the 512-dimensional cell embeddings in scGPT for cells in M.S. and Mye., respectively. *(M)* scGPT’s performance on Mye., M.S., and the hPancreas datasets.

Next, we tested the model on a disease dataset of multiple sclerosis (M.S.). The model was finetuned on a reference partition of healthy human immune cells and evaluated on the prediction for the cells with M.S. condition (Figure 2D). The fine-tuned model demonstrated strong alignment with the cell type annotations provided by the original study and achieved a high accuracy of around 0.85 (Figure 2I,J). Additionally, to investigate the impact of pre-training on fine-tuning, we separately pre-trained scGPT models on smaller subsets of the collected normal human cells, including 30K, 300K, and 3M cells, respectively. Interestingly, the results revealed a *scaling law*, where larger pre-training datasets consistently contribute to better fine-tuning performance (Figur 2E,F). This can be exemplified by the outlined regions in Figure 2E, where the fine-tuned model gradually gained the near-perfect capability of distinguishing expiatory neurons at different cortical layers as the pretraining dataset increased.

Furthermore, we applied the model to a more challenging scenario for generalization across disease types, using a tumor-infiltrating myeloid dataset (Mye.) [33]. The model was fine-tuned on six cancer types in a reference data partition (Online Methods 4) and evaluated on the query partition of three unseen cancer types (Figure 2G). The results demonstrated high precision in distinguishing immune cell subtypes (Figure 2H,K), and the cell embeddings exhibited clear separability among different cell types (Figure 2L). The predictions reached high precision (*≥* 0.7) for most cell types. Finally, we benchmarked the fine-tuned scGPT against two other recent transformer-based methods, TOSICA [34] and scBert [35], across the three datasets (Online Methods 4.7). scGPT constantly outperforms the other methods on all classification metrics, including *Accuracy, Precision, Recall*, and *MacroF* 1 (Figure 2J).

### 2.3 scGPT predicts unseen genetic perturbation responses

Recent advancements in sequencing and gene editing techniques have greatly facilitated large-scale perturbation experiments, allowing for the investigation of cellular responses to various genetic perturbations. The approach holds immense promise for uncovering novel gene interactions and advancing regenerative medicine. However, the vast combinatorial space of potential gene perturbations quickly surpasses the practical limits of experimental feasibility. To overcome this limitation, scGPT can be employed to leverage the knowledge gained from cellular responses in known experiments and extrapolate them to predict responses in unknown scenarios. The utilization of self-attention mechanisms over the gene dimension enables the encoding of intricate interactions between perturbed genes and the responses of other genes. By leveraging this capability, scGPT can effectively learn from existing experimental data and accurately predict the gene expression following perturbation.

#### Prediction of Unseen Gene Perturbations

For the perturbation prediction task, we evaluated our model using two perturbation datasets of K562 leukemia cell lines: the Adamson Pertub-seq dataset [36] consisting of 87 one-gene perturbations and the Norman Perturb-Seq dataset [37] consisting of 131 two-gene perturbations and 105 one-gene perturbations. To assess scGPT’s perturbation prediction capability, we fine-tuned the model on a subset of perturbations and tested it on perturbations involving unseen genes (Online Methods 4.5). We measured the performance using the Pearson correlation (*corr*) between the predicted and actual expression values after perturbation. We calculated the correlation scores based on the top 20 most significantly changed genes for each perturbation, namely (DE) genes. Additionally, we adopted a variant of the Pearson metric, denoted as *corr(*Δ), which measures the correlation based on the magnitude of expression change post-perturbation compared to the control. These evaluations follow the choices from GEARS [38]. See Supplementary Note S.5 for the details on metric calculations. We conducted a performance comparison between scGPT and two other recent methods, GEARS [38] and CPA [39]. Our results demonstrate that scGPT achieves the highest correlation on both datasets (Figure 3A). Additionally, we visualize the predictions for two example perturbations in the Adamson dataset in Figure 3B, where scGPT accurately predicts the trend of expression change for all top 20 DE genes.

**Figure 3:**
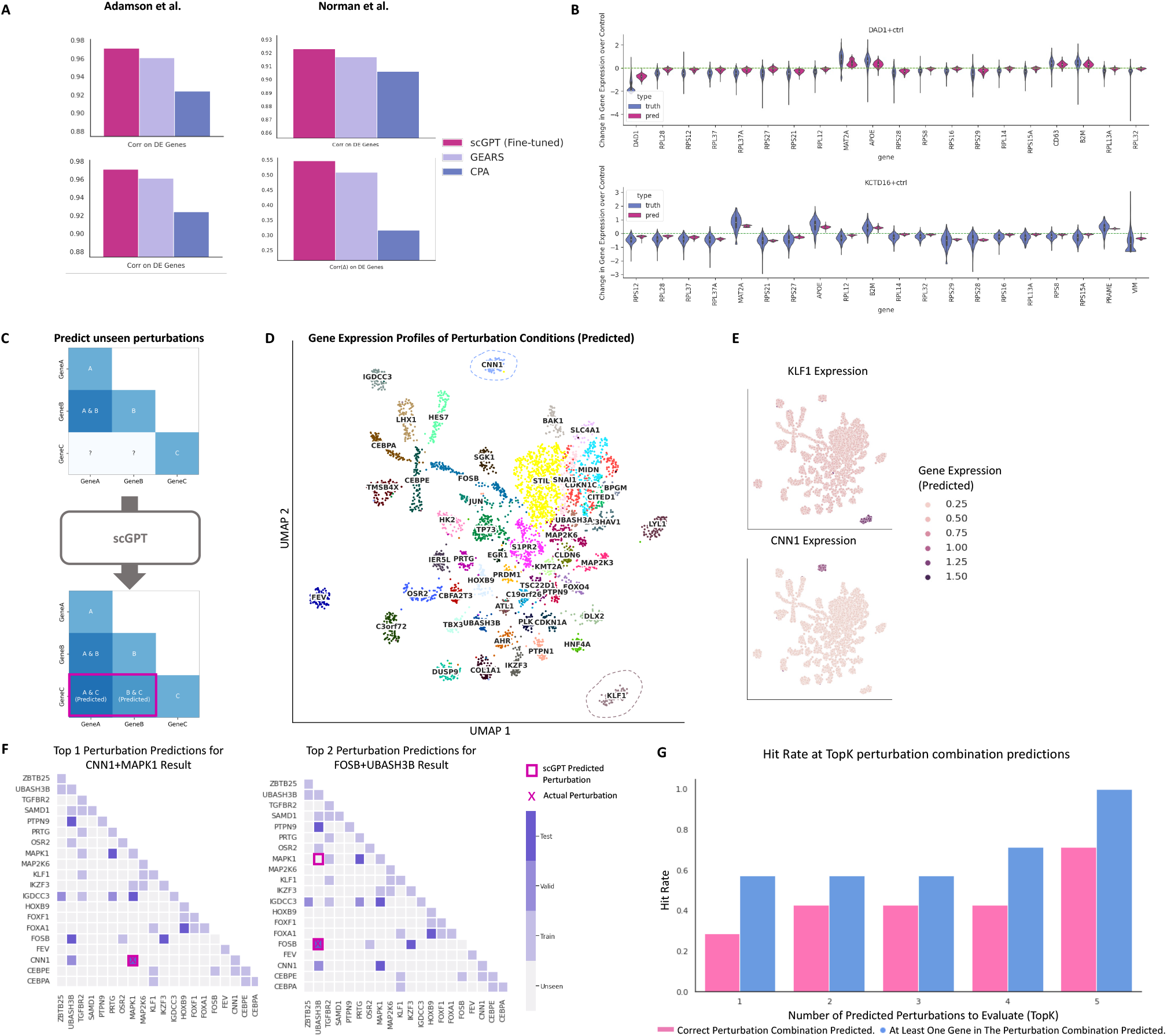
Prediction results for perturbation response and reverse perturbation. *(A)* Comparison between scGPT and other perturbation prediction methods. Two evaluation metrics are reported. *(B)* On the Adamson dataset, distribution of predicted and actual gene expression of top 20 differentially expressed genes. *(C)* Illustration diagram for predicting unseen perturbation responses using scGPT. *(D)* UMAP of predicted gene expression profiles of the perturbation conditions. The UMAP is colored by Leiden clusters and labeled by the dominant gene of each cluster. *(E)* Expression patterns of two selected perturbed genes over the UMAP of perturbation conditions. *(F)* Visualization of the possible perturbation combinations over the perturbation combination space of 20 genes. The grid is colored by experiment type (Train, Valid, Test, Unseen). All predicted perturbations are highlighted by squared boxes, and the actual source perturbation is marked by a cross. *(G)* Top 1-5 accuracy for correct and relevant predictions among the 7 test cases. The relevant predictions (blue) indicate at least one of the perturbed genes in the perturbation combination is found in the predictions.

The ability to predict unseen perturbation responses could expand the scope of perturbation experiments, as depicted in Figure 3C. To explore the expanded space of predicted perturbation responses, we conducted clustering analysis using the Norman dataset to validate the biologically relevant functional signals. The original Perturb-seq study covered 236 one-gene or two-gene perturbations targeting 105 genes. However, considering all possible combinations of these target genes, there are a total of 5,565 potential perturbations. This means that the experimental Perturb-seq only represents 5% of the entire perturbation space. Therefore, we applied the fine-tuned scGPT to expand the perturbation *in-silico* and visualized the predicted mean response for each perturbation in Figure 3D using UMAP. Using the annotations from the original study, we found the perturbation conditions of the same functional groups gathered at nearby regions (Supplementary Figure S4). Next, we clustered the predicted expression using Leiden [40] and mapped each data point with the corresponding perturbed genes. Interestingly, we observed that the clusters exhibited a high association with the “dominant gene” within the perturbation combinations. For example, the circled cluster associated with the *KLF1* gene indicated that the data points in this cluster underwent combined perturbations involving *KLF1* along with another gene (i.e., *KLF1 + X*). Using *KLF1* and *CNN1* clusters as two examples, we further validated that the corresponding predicted expression was exclusively high in these regions (Figure 3E), which aligns with expected outcomes of the CRIPRa Perturb-seq experiments in the Norman dataset. These dominant gene clusters demonstrate the capability of scGPT to uncover associations between perturbation combinations, providing insights into their combined effects on gene expression profiles.

#### In-silico *Reverse Perturbation* Prediction

scGPT is also capable of predicting the source of genetic perturbation for a given resulting cell state, which we refer to as in-silico *reverse perturbation* prediction. An ideal prediction model conducting such reverse prediction can be used to infer important driving genes for lineage development or facilitate the discovery of potential therapeutic gene targets. A hypothetical example application of such capability could be to predict CRISPR target genes that influence cells to recover from a disease state. To showcase the effectiveness of reverse perturbation prediction, we utilized a subset of the Norman dataset focusing on perturbations involving 20 genes (Figure 3F). This combinatorial space consists of a total of 210 one-gene or two-gene perturbation combinations. We fine-tuned scGPT using 39 (19%) known perturbations (the train group in Figure 3F). We then tested the model on queries of unseen perturbed cell states, and scGPT successfully predicted the source of perturbations that would generate the observed results. Specifically, after ranking the top predictions for seven test examples, we found that scGPT accurately retrieved the actual source perturbations. For example, scGPT ranked the correct perturbation of *CNN1+MAPK1* as the top prediction for one test example, and the correct perturbation of *FOSB+UBASH3B* was ranked as the second prediction for another case (Figure 3F). Overall, scGPT identified 71.4% correct perturbations (5/7 test cases) within the top 5 predictions (the red bars in Figure 3G). We envision these predictions can be used for planning perturbation experiments by maximizing the possibility of deriving target cell states. Compared to random tryouts, which would on average require 105.5 attempts out of the 210 possible perturbations in this subset, finding the correct source of genetic change within 5 attempts represents a 95.2% speedup. The reverse perturbation predictions generated by scGPT offer a valuable tool for accelerating the discovery of important genetic drivers and optimizing perturbation experiments.

### 2.4 scGPT enables accurate multi-batch and multi-omics integration

#### Multi-Batch scRNA-seq Integration

Integrating multiple scRNA-seq data from different batches poses unique challenges in simultaneously preserving the biological variance of integrated datasets and removing technical batch effects. In our benchmarking experiments, we compared scGPT with three popular integration methods: scVI [41], Seurat [42], and Harmony [43]. The evaluation was conducted on three integration datasets, namely the COVID-19 (18 batches) [13], PBMC 10K (2 batches) [44], and Perirhinal Cortex (2 batches) [45]. In the PBMC 10K dataset, scGPT stands out as the only method that successfully separates all the cell types. The other methods struggle with accurately distinguishing CD14+ Monocytes and CD8 T cells from other cell types (Figure 4A). This superior integration performance of scGPT is further supported by its high biological conservation score, with an *AvgBIO* score of 0.821, which is 5-10% higher than the compared methods. The *AvgBIO* score aggregates three cell type clustering metrics *NMI*_*cell*_, *ARI*_*cell*_, and *ASW*_*cell*_, as detailed in Supplementary Online Methods S.5. Notably, scGPT also demonstrates a considerable performance for integrating PBMC 10K dataset even without fine-tuning (Supplementary Figure S5), highlighting the generalizability of the pretraining. In the context of the Perirhinal Cortex dataset, scGPT remains competitive against all other methodologies (Supplementary Figure S6C). This finding highlights the transferability and robustness of the features learned from the whole-human dataset, even when applied to specific organs/tissues such as the brain. Furthermore, in Supplementary Figure S6, scGPT consistently achieves competitive scores across all integration metrics and demonstrates strong conservation of biological signals. In terms of overall performance, considering both biological conservation and batch correction, scGPT ranks at the top (See detailed metrics in Supplementary Table S2 and Supplementary Figure S7). Additionally, we have developed strategies to accelerate the fine-tuning process for the integration task, including freezing specific model layers and excluding genes with no expression, while maintaining comparable results to our original approach (Supplementary Note S.3).

**Figure 4:**
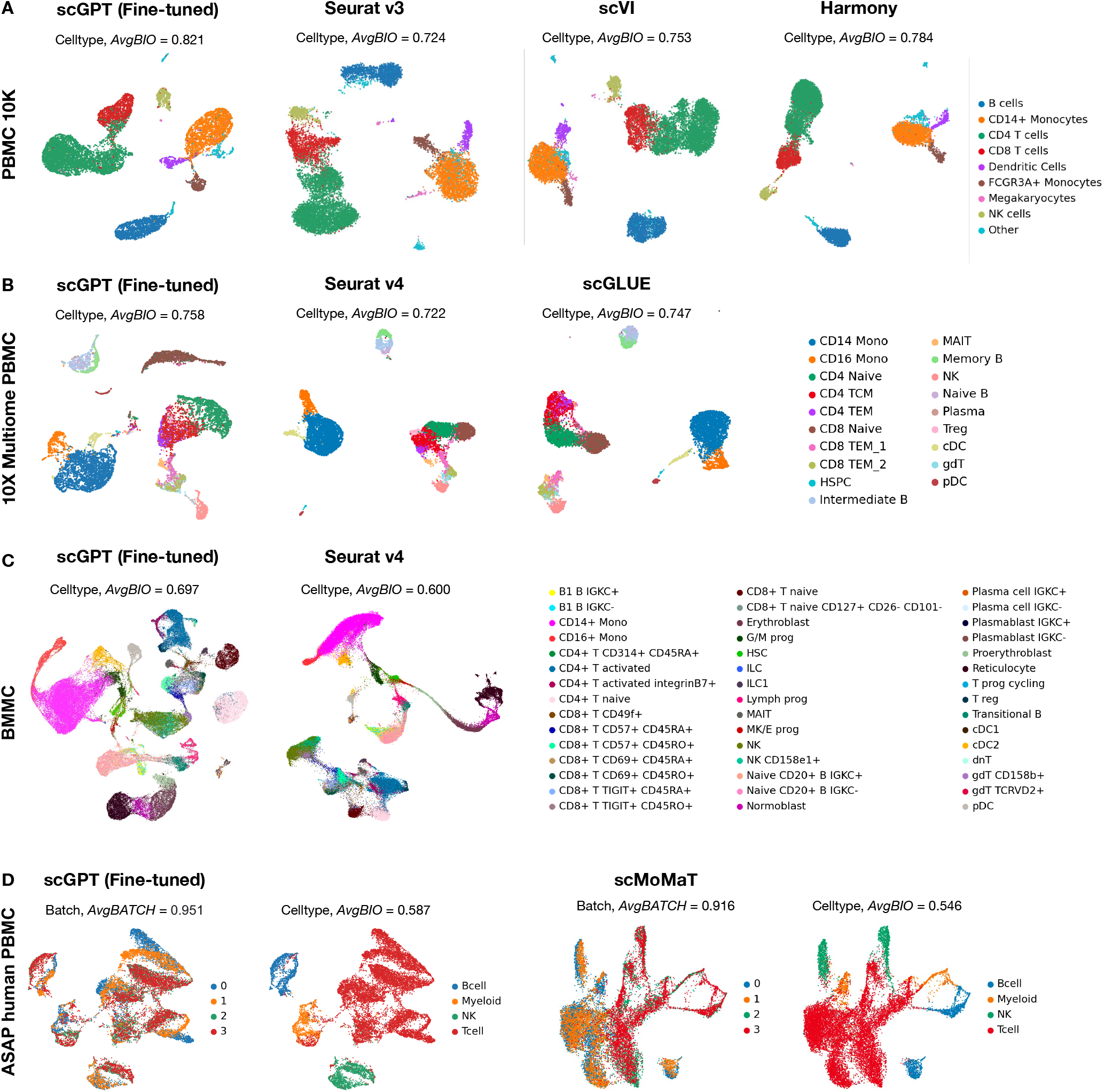
*(A)* Benchmark of the fine-tuned scGPT on the PBMC 10K dataset for cell type clustering task. The UMAP plot of learned cell embeddings was colored by *cell types. (B)* Benchmark of the fine-tuned scGPT model with scGLUE[14] and Seurat v4[46] on the 10x Multiome PBMC [47] dataset (paired RNA+ATAC data) for cell type clustering task. *(C)* Benchmark of the finetuned scGPT model with Seurat v4[46] on the BMMC [48] dataset (paired RNA+Protein data) for cell type clustering task. *(D)* Benchmark of scGPT with scMoMat[15] on the ASAP PBMC [49] dataset (mosaic RNA+ATAC+Protein data) for batch correction and cell type clustering tasks. The UMAP plots of learned gene embeddings were colored by *sequencing batches* (left) and *cell types* (right).

#### Single-Cell Multi-Omic Integration

Single-cell multi-omic (scMultiomic) data, which combines multiple views of genetic regulation such as epigenetic, transcriptomic, and translation activities, presents a unique challenge in aggregating cell representations while preserving biological signals [9, 10]. scGPT addresses this challenge by effectively extracting integrated cell embeddings across different omics. In the case of the 10X Multiome PBMC dataset [47], which includes joint gene expression and chromatin accessibility measurements, we compared scGPT with two state-of-the-art methods, scGLUE [14] and Seurat v4 [46], in terms of cell type integration performance. As depicted in Figure 4B, scGPT stands out as the only method that successfully generates a distinct cluster for CD8 Naive cells, while the other two methods fail to do so. Next, we tested scGPT on the paired gene expression and protein abundance dataset BMMC [48] as illustrated in Figure 4C. This dataset contains additional complexity from the large data size (90 thousand cells), multiple batches (12 donors), and fine-grained subgroup annotations (48 cell types). scGPT presented more defined cluster structures compared to Seurat v4, with a 9% improvement in the *AvgBIO* score. Notably, scGPT was able to separate CD4+ T naive and CD4+ T activated cells as two distinct clusters. It also teased apart integrinB7+ activated CD4+ T cells from other CD4+ T cells, which further endorsed the model’s ability to capture the subtle differences between immune cell subgroups. Overall, scGPT demonstrates superior cell-type clustering performance in the paired data setting and exhibits robustness across diverse biological conservation metrics benchmarked (Figure 4D and Supplementary Table S3).

scGPT excels in integrating multi-modal batches from mosaic scMultiomic data through joint representation learning. In the mosaic data integration setting, sequenced samples share some, but not all, data modalities, posing a challenge for integration methods. To showcase the capabilities of scGPT in this context, we utilized the ASAP human PBMC dataset [49] as an example. This dataset consists of four sequencing batches with three data modalities: gene expression and protein abundance data from CITE-seq in the first two batches, and chromatin accessibility and protein abundance measurements from ASAP-seq in the second two batches. In the benchmark experiment with scMoMat[15], scGPT demonstrates superior batch correction performance as shown in Figure 4D, especially in the rarer cell groups B, Myeloid, and NK cells. In comparison, scMoMat produced two distinct clusters for each cell type corresponding to the first two and second two batches, indicating failure to learn a joint representation. scGPT achieves a higher overall batch correction score *AvgBATCH* of 0.951 compared to scMoMat (*AvgBATCH* = 0.916). scGPT’s biological conservation metrics also compare favorably to scMoMat’s, which further indicates the robustness of multi-modal batch correction without interfering with the biological signals (Figure 4D and Supplementary Table S3).

### 2.5 scGPT uncovers gene regulatory networks for specific cell states and perturbation conditions

The interactivity between transcription factors, cofactors, enhancers, and target genes underlying a Gene Regulatory Network (GRN) mediates important biological processes. Existing GRN inference methods often rely on correlation in static gene expression or pseudo-time estimates as a proxy for causal graphs [50]. scGPT, optimized by generative modelling of gene expression, implicitly encodes such relationships in its gene embeddings as well as attention maps. The gene embeddings construct a similarity network that entails gene-gene interactions on the dataset level. The attention maps further capture the unique gene network activation patterns across different cell states. In this study, we validate the gene network extracted by scGPT against known biology and explore its applicability to gene program discovery.

scGPT demonstrates its ability to group functionally related genes and differentiate functionally distinct genes via the learned gene token embeddings. In Figure 5A, we conducted a sanity check by visualizing the similarity network of the human leukocyte antigen (HLA) proteins using the pre-trained gene embeddings from the pre-trained scGPT model. In this zero-shot setting, the scGPT model successfully highlights two clusters corresponding to the well-characterized HLA classes: HLA class I and HLA class II genes. These classes encode antigen-presenting proteins that play different roles in immune contexts. For example, HLA class I proteins (such as *HLA-A*, -*C*, and -*E*) are recognized by CD8+ T cells and mediate cytotoxic effects, while HLA class II proteins (such as HLA-DR, -DP, and -DQ) are recognized by CD4+ T cells and trigger broader helper functions [51]. In addition, we fine-tuned the scGPT model on the Immune Human dataset and explored the CD gene network specific to the immune cell types present in this dataset. We employed the same fine-tuning strategy as used for the integration task (Online Methods 4.5) for the purpose of GRN analysis. The pre-trained scGPT model successfully identifies the group of genes (*CD3E, D*, and *G*) encoding the T3 complex for T-cell activation, as well as *CD79A* and *CD79B* for B-cell signaling, and *CD8A* and *CD8B* as co-receptors for HLA class I molecules [52] (Figure 5B). Furthermore, the fine-tuned scGPT model highlights the connection between *CD36* and *CD14*, which serve as markers for monocytes and macrophages (Figure 5B). These findings demonstrate scGPT’s ability to generalize the knowledge learned during pre-training and extract information relevant to specific tissues or cell types after fine-tuning.

**Figure 5:**
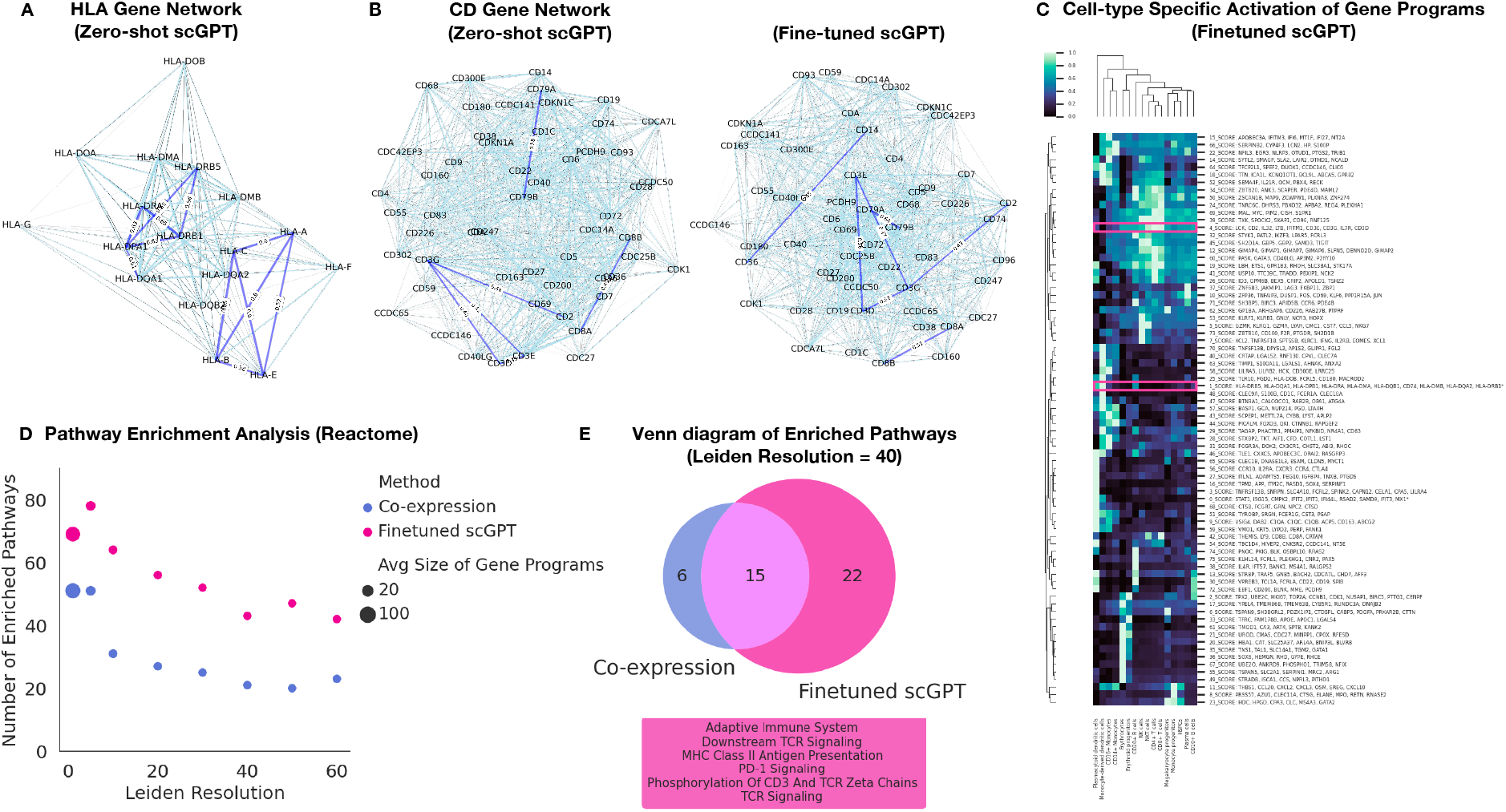
*(A)* HLA gene network from zero-shot scGPT. *(B)* CD gene network from the zero-shot (i.e., pre-trained) and fine-tuned scGPT on the Immune Human dataset. *(C)* Cell-type specific activation among scGPT-extracted gene programs in the Immune Human dataset. *(D)* Pathway enrichment analysis of the gene programs extracted by scGPT and co-expression network in the Immune Human dataset. The number of enriched pathways from scGPT’s gene programs was compared with the co-expression method at different Leiden resolutions. *(E)* Venn diagram to compare the overlap and differences between the enriched pathways identified by co-expression versus scGPT. Some example pathways unique to scGPT and specific to the adaptive immune functions are highlighted in the text box.

scGPT is able to uncover meaningful gene programs that exhibit cell-type specific activation via purely unsupervised pretraining and fine-tuning. At the inference stage, gene programs are subsequently selected and clustered using the gene embeddings from scGPT (Online Methods 4.5). In Figure 5C, we visualized the gene programs extracted by the fine-tuned scGPT model on the highly variable genes in the Immune Human dataset [53], and their expression in different cell types. We observed that a set of HLA class II genes was identified as group 1. Similarly, the CD3 genes involved in the T3 complex were identified as group 4, with the highest expression present in T cells. These findings confirm that scGPT’s inferred gene programs capture biologically meaningful functional groups.

In order to systematically validate the extracted gene programs, we performed pathway enrichment analysis against the Reactome database [54] and identified high-confidence “pathway hits” using stringent multiple-testing correction [55] (Online Methods 4.7). In Figure 5D, we compare the results obtained from scGPT with those from the co-expression network. Notably, scGPT consistently demonstrates a significantly higher number of enriched pathways across all clustering resolutions. Furthermore, we examined the similarities and differences in the identified pathways between scGPT and the co-expression network, as depicted in Figure 5E. Both methods identified 15 common pathways, including those associated with the cell cycle and the immune system. However, scGPT uniquely identified additional 22 pathways, out of which 14 were immune-related. Notably, scGPT specifically highlighted pathways related to the adaptive immune system, T-cell receptor (TCR) signaling, PD-1 signaling, and MHC II presentation, which were not captured by the co-expression network. This is concordant with the fact that adaptive immune populations exist in the fine-tuning datasets. These findings demonstrate scGPT’s superior ability to capture intricate gene-gene connections and unravel specific mechanisms within a broader biological context. The detailed list of enriched pathways is provided in Supplementary Table S4.

In addition to dataset-level gene network inference using gene embeddings, scGPT’s attention mechanism enables it to capture gene-gene interactions at the single-cell level. scGPT extracts cellstate-specific network activations by aggregating single-cell signals from the attention maps. This provides insights into the context-specific gene regulatory interactions within individual cells, which may vary across different cell states and conditions. For example, in a perturbation experiment, scGPT examines the changes in gene network activation pre- and post-perturbation to infer which genes are most influenced by each perturbed gene (Figure 6A and Online Methods 4.5). In the Adamson CRISPR interference dataset [36], scGPT identified the top 20 genes most influenced by *DDIT3* transcription factor repression, which are all found to be signaling targets of *DDIT3* in the Chip-Atlas database [56] (Figure 6B). Moreover, scGPT captured distinct pathway activation patterns among the top 100 most influenced genes by *DDIT3* in the control versus *DDIT3* -knockout settings. Notably, the ATF6 pathway identified in *DDIT3* -knockout setting is known to mediate unfolded protein response and regulates cell apoptosis [57, 58]. Similarly, in the case of *BHLHE40* repression, 19 out of the top 20 most influenced genes are found to be ChIP-seq predicted targets of this transcription factor (Figure 6C). The pathway activation profile highlighting DNA synthesis and mitosis reflects the role of *BHLHE40* in cell cycle regulation [59]. These attention-based findings further validated scGPT’s learned gene network on a cell-state level, providing additional interpretability to the model’s learned biology.

**Figure 6:**
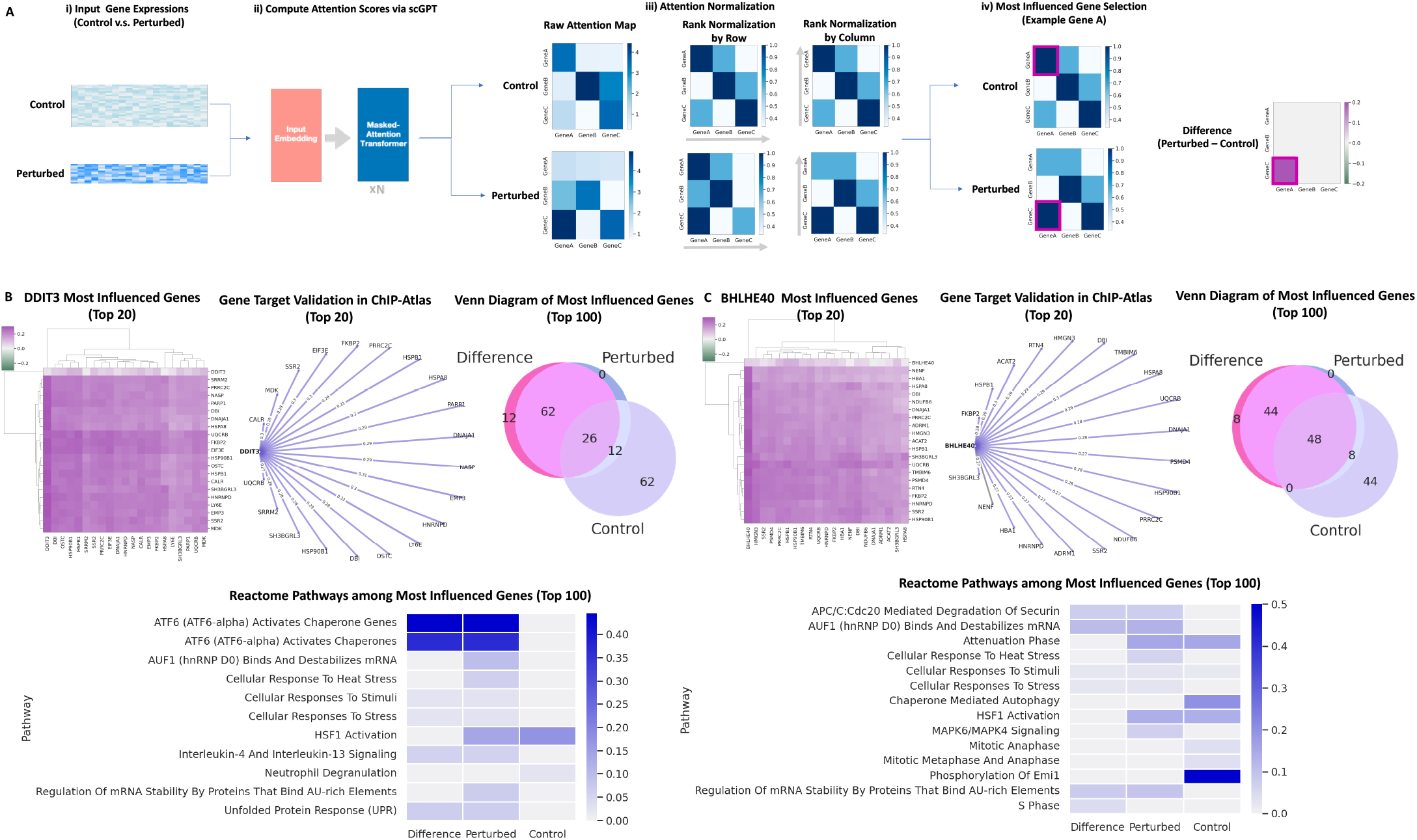
*(A)* Attention-based GRN discovery workflow for perturbation data. Attention scores in control and perturbed cell states are obtained and ranked by row and column consecutively. Most influenced genes in *Control, Perturbed*, and the *Difference* settings are selected accordingly. *(B)* GRN analysis for DDIT3 transcription factor repression. The gene connectivity heatmap presents post-perturbation changes in the network of the top 20 genes most influenced by DDIT3 repression. The gene target network graph showcases the top 20 genes validated in the CHIP-Atlas database, with ChIP-seq predicted targets highlighted in purple. The Venn diagram compares the overlaps and differences among the top 100 most influenced gene sets identified in three selection settings (i.e., *Control, Perturbed*, and post-perturbation *Difference*). The pathway heatmap showcases the difference in Reactome pathways identified from the top 100 most influenced genes across these three selection settings. The color indicates how strong each pathway is represented by the percentage of gene overlap. *(C)* GRN analysis for BHLHE40 transcription factor repression, visualized in a similar manner.

These aforementioned results showcase the potential of scGPT for exploring gene-gene interactions in a general setting and for specific cell types and perturbation conditions. More specifically, we demonstrate its ability to perform unsupervised gene program discovery on new datasets that disentangle fine-grained biological mechanisms. The attention map further provides a prediction-based importance ranking approach that identifies cell-state-specific gene targets taking part in dynamic processes.

### 2.6 Analyzing the transfer learning of pretrained knowledge: the scaling law and in-context effects

In previous sections, scGPT has demonstrated great potential via fine-tuning in a transfer learning manner. We further confirmed the benefits of using the foundation model by comparing it with similar transformer models trained for each downstream task from scratch without pretraining (denoted as scGPT (from-scratch)). The results are listed in Supplementary Tables S1, S2, S3, where the fine-tuned scGPT consistently showcases performance gain for tasks such as integration and cell type annotation. Given the observed contributions of the foundation model for downstream tasks, we are further interested in exploring the factors influencing the transfer learning process.

Firstly, we delve into the relationship between *pre-training data size* and the performance of fine-tuned models. We pre-trained a series of scGPT models of the same number of parameters but using different sizes of data, from 30K to 33M sequenced normal human cells. Figure 2D and E illustrate the actual performance of fine-tuning for the cell type annotation task using these various pre-trained models. We observed the performance of fine-tuned models improved as the size of the pre-training data increased. These results confirm the *scaling law*, suggesting that larger pre-training data size leads to better pre-trained embeddings and improved performance on downstream tasks. On the other hand, we acknowledge the importance of considering model size alongside data size. Previous research has shown a correlation between optimal model size and data size, as discussed [60]. While the scaling law trend in Figure 2E may appear to reach a saturation point after the 3M data size, we hypothesize that increasing the model size would lead to improved performance even at 33M data size and beyond. We leave the exploration of larger model sizes to future work.

Notably, our findings also align with the scaling law reported in natural language models [60], highlighting the significant role of data size in model performance. The crucial role of pretraining data size in fine-tuning results points to a promising future for pretrained models in the single-cell domain. As larger and more diverse datasets become available, we can anticipate further improvements in model performance, advancing our understanding of cellular processes. Pretrained models will continue to expand their capabilities, facilitating breakthroughs in single-cell research and unlocking new discoveries in cellular biology.

The second factor we explored is the influence of context-specific pre-training. Here, an *incontext usage* refers to an scGPT model that is pre-trained on specific cell types and then fine-tuned for a downstream task on similar cell types. To explore the influence of this factor, we pre-trained seven organ-specific models on the normal human cells from individual major organs (Figure 1D) and another model for pan-cancer cells. We verified the efficacy of the pretraining by visualizing the cell embeddings of the pre-training data: the pan-cancer model cell embeddings accurately separate different cancer types (Supplementary Figure S2). The organ-specific models are able to reveal cell heterogeneity of corresponding organs (Supplementary Figure S3). Interestingly, since the training data per organ were collected in different sizes, the organ-specific pretrained models again confirmed the scaling law: models pre-trained on larger data (*>* 800, 000 cells) exhibited cell embeddings better separating the cell types, as shown in Supplementary Figure S3. Next, removing the influence of data sizes, we used the rest five organ-specific models (i.e., the brain, blood, heart, lung, and kidney models) with sufficient training data for the analysis of in-context effects. The COVID-19 dataset serves as an ideal task for this investigation, as it comprises diverse cellular compositions. Our analysis reveals a clear correlation between the relevance of the model’s context in pre-training and its subsequent performance for integrating the COVID-19 dataset (Supplementary Figure S8). The top performers in data integration tasks were models pretrained on whole-human, blood, and lung datasets, which closely aligned with the cell types present in the COVID-19 dataset. Notably, even a brain pre-trained model, despite being trained on a substantial dataset of 13 million cells, trailed in performance by 8% compared to the blood pre-trained model with similar dataset size. This emphasizes the importance of aligning the cellular context in pretraining with the target dataset for superior results in downstream tasks. While considering the cellular context is essential, the whole-human pre-trained model emerges as a versatile and reliable option for a wide range of applications.

In summary, our analysis demonstrates the contribution of pretrained knowledge to downstream analysis, the scaling law between pre-training data size and fine-tuning results, and the impact of context-specific pre-training. These findings highlight the potential of pretrained models to advance our understanding of cellular biology and pave the way for future discoveries.

## 3 Discussion

We introduce scGPT, a pioneering generative pretrained foundation model that harnesses the power of pre-trained transformers on a vast amount of single-cell data. Building upon the success of self-supervised pre-training in language models like chatGPT and GPT4 [18, 19], we adopted a similar approach in the single-cell domain to unravel complex biological interactions. The use of transformers in scGPT enables the simultaneous learning of gene and cell embeddings, which facilitates the modeling of various aspects of cellular processes. By leveraging the attention mechanism of transformers, scGPT captures gene-to-gene interactions at the single-cell level, providing an additional layer of interpretability.

We demonstrated the benefits of pre-training with comprehensive experiments in both zero-shot and fine-tuning settings. The pre-trained model itself is a universal feature extractor. It showcases strong capabilities of extrapolating to unseen datasets, presenting meaningful cell clustering in zeroshot experiments. In addition, the learned gene networks in scGPT exhibit strong alignment with known functional connections. The model’s ability to accurately capture gene-gene interactions and reflect known biological knowledge provides confidence in its capacity to uncover meaningful insights in single-cell biology. Furthermore, the pre-trained model’s knowledge can be transferred to multiple downstream tasks through fine-tuning. In tasks such as cell type annotation, multibatch and multi-omic integration, the fine-tuned scGPT model consistently outperforms models trained from scratch. This demonstrates the valuable contribution of the pre-trained model to downstream tasks, enabling more accurate and biologically meaningful analyses. Overall, our experiments highlight the efficacy of pre-training in scGPT, showcasing its ability to generalize, capture gene networks, and improve performance in downstream tasks through transfer learning.

For future directions, we plan to pre-train on a larger-scale dataset with more diversity, including multi-omic data, spatial omics, and various diseased conditions. It is also interesting to incorporate perturbation and temporal data in the pre-training stage, enabling the model to learn causal relationships and infer how genes and cells respond to changes over time. More importantly, we would like to validate the pre-trained model on a wider range of biologically meaningful tasks to understand and interpret what the pre-trained model has learned. We also aim to explore in-context instruction learning for single-cell data. This involves developing techniques that allow the pre-trained model to understand and adapt to different tasks and contexts in a zero-shot setting, without the need for fine-tuning. By enabling scGPT to grasp the nuances and specific requirements of different analyses, we can enhance its usability and applicability in a wide range of research scenarios. We envision that the pre-training paradigm be readily integrated into singlecell research, and serve as a foundation to leverage the existing knowledge from the exponentially growing cell atlases for new discoveries.

## 4 Methods

### 4.1 Input embeddings

The single-cell sequencing data is processed into a cell-by-gene matrix, ***X*** *∈* ℝ^*N×G*^, where each element *X*_*i,j*_ *∈* ℝ^+^ represents the read count of an RNA molecule in the case of scRNA-seq or a peak region in the case of scATAC-seq. Specifically, in scRNA-seq, the element denotes the RNA abundance for gene *j ∈* 0, 1, …, *G* in cell *i ∈* 0, 1, …, *N*. In subsequent sections, we will refer to this matrix as the raw matrix. The input to scGPT consists of three main components: (1) gene (or peak) tokens, (2) expression values, and (3) condition tokens. For each modelling task, the gene tokens and expression values are pre-processed from the raw count matrix ***X*** accordingly.

#### Gene Tokens

In the scGPT framework, each gene is considered the smallest unit of information, analogous to a word in natural language generation (NLG). We therefore use gene names as tokens, and assign each gene *g*_*j*_ a unique integer identifier *id*(*g*_*j*_). These identifiers form the vocabulary of tokens used in scGPT. This approach offers great flexibility to harmonize multiple studies with different gene sets (i.e., generated by distinct sequencing technologies or pre-processing pipelines). Specifically, different sets of gene tokens can be integrated into a common vocabulary by taking the union set of all genes across studies. Additionally, we incorporate special tokens in the vocabulary, such as *< cls >* for aggregating all genes into a cell representation, and *< pad >* for padding the input to a fixed length. Conceptually, we draw parallels between gene tokens and word tokens in NLG. The input gene tokens of each cell *i* are hence represented by a vector 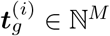 :

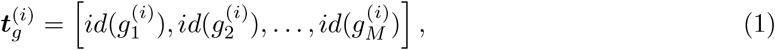

where *M* is a pre-defined maximum input length.

#### Expression Values

The gene expression matrix *X* requires additional processing before being used as input for modelling. A fundamental challenge in gene expression modelling is the variability in absolute magnitudes across different sequencing protocols [61]. Variations in sequencing depths and the presence of sparsely expressed genes result in significant differences in data scales among different batches of sequencing samples. These differences are not easily mitigated with common preprocessing techniques such as transcripts per million (TPM) normalization and *log*1*p* transformation [62]. Even after these transformations, the same absolute value can convey different “semantic” meanings across sequencing batches. To address this scale difference, we propose the ***value binning*** technique to convert all expression counts into relative values. For each non-zero expression count in each cell, we calculate the raw absolute values and divide them into *B* consecutive intervals [*b*_*k*_, *b*_*k+1*_], where *k ∈ {*1, 2, …, *B}*. Each interval represents an equal portion of all expressed genes (1*/B*). It is important to note that a new set of bin edges is computed for each cell, so the interval edges *b*_*k*_ may vary among cells. The binned expression value 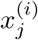 for cell *i* is defined as:

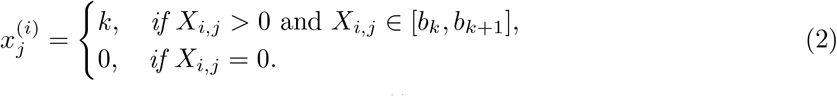

Through this binning technique, the semantic meaning of 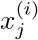 is consistent across cells from various sequencing batches. For instance, a value of 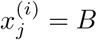 consistently indicates the highest expression among genes. Notably for fine-tuning tasks, we also performed *log*1*p* transformation and highly variable gene selection [63] before the value binning step. To simplify the notation, we use *X*_*i,j*_ to represent both the raw and preprocessed data matrices prior to binning. Therefore, the final input vector of binned expression values for cell *i* is denoted as

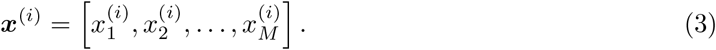

#### Condition Tokens

The condition tokens encompass diverse meta information associated with individual genes, such as perturbation experiment alterations (indicated by perturbation tokens). To represent position-wise condition tokens, we utilize an input vector that shares the same dimension as the input genes. This vector is denoted as:

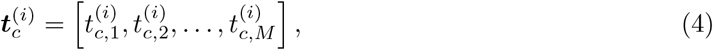

where 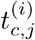 represents an integer index corresponding to a condition.

#### Embedding layers

We utilize the conventional embedding layers^1^ emb_g_ and emb_c_ for the gene tokens and condition tokens, respectively, to facilitate the mapping of each token to a fixed-length embedding vector of dimension *D*. We employ fully connected layers, denoted as emb_x_, for the binned expression values to enhance expressivity. This choice enables the modelling of the ordinal relation of gene expression values. Consequently, the final embedding *h*^(*i*)^ *∈* ℝ^*M×D*^ for cell *i* is defined as,

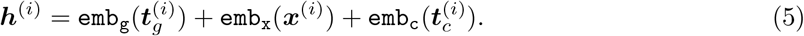

### 4.2 Cell and gene expression modelling by transformers

#### 4.2.1 scGPT Transformer

We employ the self-attention transformer [27, 64] to encode the complete input embedding ***h***^(*i*)^ in equation 5. The self-attention mechanism operates on the sequence of *M* embedding vectors, making it particularly suitable for capturing interactions between genes. The output of the stacked transformer blocks can be defined as follows:

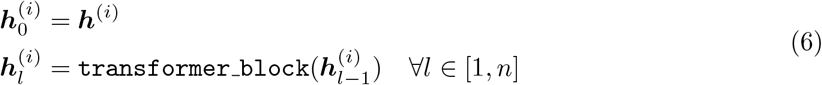

We utilize the resulting representation 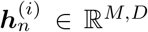 for both gene-level and cell-level tasks. Gene-level fine-tuning objectives (Online Methods 4.4) are directly applied. Examples include the gene expression prediction (GEP) objective and the perturbed expression prediction task (perturb-GEP). For cell-level tasks, we first integrate 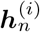 into a cell embedding vector (Online Methods 4.2.2). An example would be the cell type assignment task, where the cell embeddings are used to predict cell type labels by an added classifier in the CLS training objective.

The input dimension *M* can reach tens of thousands of genes, significantly exceeding the input length of conventional transformers commonly used in NLG. To address this challenge and ensure efficient self-attention mechanisms, we leverage the accelerated self-attention implementation by Flash-Attention [65]. This implementation effectively enhances the model capacity and enables the effective processing of large input dimensions. Although Flash-Attention was adopted, any efficient transformers can be potentially utilized for scGPT as well, such as Transformers with linear complexity (Linformer) [66] and Kernelized Self-Attention (KSA) [67].

#### 4.2.2 Cell representation

Each cell is analogous to a “sentence” composed of genes, and its representation 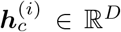 is obtained by aggregating the learned gene-level representations 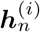. Various pooling operations, such as element-wise mean-pooling or weighted-pooling, can be readily employed in this context. In this study, we opt to employ a special token *< cls >* for the cell representation, enabling the model to learn the pooling operation within transformer blocks. The *< cls >* token is appended to the beginning of the input tokens, and the final embedding at this position is extracted as the cell representation. Consequently, the cell embedding ***h***^(*i*)^ can be extracted by the corresponding row in the stacked final-layer embeddings 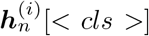, where the [*< cls >*] operation retrieves the row at the index of the *< cls >* token position.

#### 4.2.3 Representation for batch and modality

We use additional sets of tokens to represent different sequencing batches and sequencing modalities (genes from RNA-seq, peaks from ATAC-seq, etc.), specifically for the scRNA-seq and scMultiomic integration tasks. This is similar to condition tokens introduced in Online Methods 4.1, and implemented similarly using the standard embedding layers. The modality tokens 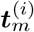 are associated with individual input features *g*_*j*_ (e.g., to indicate whether it is a gene, region or protein). The batch tokens are on the cell level originally but can be propagated to all features of a single cell as well. In other words, the same batch token 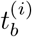 can be repeated up to the length *M* of input features of single cell *i*:

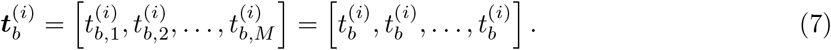

The difference between the tokens described in Online Methods 4.1 and the batch and modality tokens is that these embeddings of batch and modality tokens are not used as input to the transformer blocks. Instead, they are concatenated with the transformer output on either feature or cell level prior to entering specific fine-tuning objectives. This is to prevent the transformer from amplifying the attention within features of the same modalities while underestimating those of different modalities. Furthermore, knowing the modality and/or batch identities facilitates gene expression modelling in the downstream fine-tuning objectives. As the model learns to predict expression values conditioned on modality and/or batch identities, such biases are implicitly removed from the gene and cell representations themselves. This serves as a technique to facilitate batch correction.

As an example, in the scMultiomic integration task, we concatenate the transformer output with the sum of batch and modality embeddings. This serves as input to the downstream fine-tuning objectives for expression modelling:

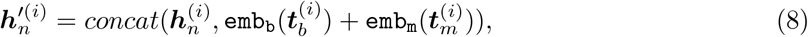

where emb_b_ and emb_m_ denote the batch and modality embedding layers respectively. 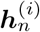 denotes the output of the transformer layer (Online Methods 4.2.1).

Alternatively, in the scRNA-seq integration task, concatenation of batch embedding with the cell representation yields the following representation as input:

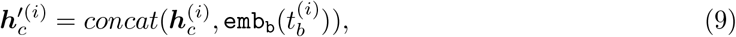

where 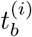 denotes the batch identity of cell *i*. 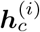 is the original cell representation in Online Methods 4.2.2.

### 4.3 Generative pre-training

#### 4.3.1 Foundation model pre-training

The foundation model is designed to be a generalizable feature extractor that can benefit a diverse range of downstream tasks. The token vocabulary used in pre-training contains the entire set of genes in the human genome. The expression values were binned prior to model pre-training (Online Methods 4.1). To speed up the training, we restrict the input to only genes with nonzero expression for each input cell. To efficiently train the model to capture gene-gene relation and gene-cell relation, we introduce a new generative training strategy with specialized attention masks as described in the following section.

#### 4.3.2 Attention mask for generative pre-training

Self-attention has been widely used to capture the co-occurrence patterns among tokens. In natural language processing, this has been achieved mainly in two ways: (1) masked token prediction used in transformer encoder models such as BERT [64] and Roberta [68], where randomly masked tokens in the input sequence are predicted in the model’s output; (2) auto-regressive generation with sequential prediction in causal transformer decoder models such as the OpenAI GPT series [69, 70, 71, 19]. The generative pre-training used in OpenAI GPT3 [71] and GPT4 [19] employs a unified framework in which the model predicts the most likely next token from a “prompt” consisting of known input tokens. This framework offers great flexibility to be utilized in various natural language generation (NLG) applications and demonstrates new capabilities such as contextual awareness in zero-shot and fine-tuned settings [72]. We believe that generative training can be beneficial to single-cell models in a similar manner. Specifically, we are interested in two tasks: (1) generating unknown gene expression values based on known gene expression, i.e., generation by “gene prompts”, and (2) generating whole genome expression given an input cell type condition, i.e., generation by “cell prompts”.

Despite similar ideas of tokens and prompts, modelling genetic reads is inherently different from natural language due to the non-sequential nature of the data. Unlike words in a sentence, the order of genes within a cell is interchangeable, and there is no equivalent concept of the “next gene” to predict. This makes it challenging to apply the causal masking formulation from GPT models directly in single-cell data. To address this challenge, we developed a specialized attention masking mechanism for scGPT that defines the order of prediction based on attention scores.

The scGPT’s attention mask supports both gene-prompt and cell-prompt generations in a unified way. The binary attention mask is applied on the self-attention map in the transformer blocks. For an input 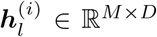 of *M* tokens (Online Methods 4.2.1), the transformer block will generate *M* query and key vectors to compute the attention map, ***A*** *∈* ℝ^*M×M*^. The attention mask is of the same size *M × M*. We visualize the attention mask in Supplementary Figure S1A, where queries are organized in rows and keys in columns. The token identity associated with each column of the mask is annotated at the bottom of the figure, namely *< cls >*, known genes, and unknown genes. Each token in the input embedding 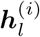 can be one of these three groups: (1) the reserved *< cls >* token for cell embedding (introduced in Online Methods 4.2.2), (2) known genes with token embeddings and expression value embeddings, and (3) unknown genes whose expression values are to be predicted. The rule of thumb for scGPT’s attention-masking is to only allow attention computation between embeddings of the “known genes” and the query gene itself. In each generation iteration, scGPT predicts the gene expression values of a new set of genes, and these genes in turn become the “known genes” in the next iteration for attention computation. This approach reflects the casual masking design with next token prediction in the conventional transformer decoders by making sequential predictions in the non-sequential single-cell data.

As illustrated in Supplementary Figure S1A, during training, we randomly pick a proportion of the genes as unknown so their expression values are omitted in the input. Attention is only applied between the known genes and the query unknown gene itself, but not onto the positions of other unknown genes. For example, the last gene to predict at position M has attention scores with the cell embedding, known genes, and itself only, but not the other unknown genes, as illustrated in the last row of the attention mask. The scGPT model predicts the expression for these unknown genes via the stacked transformer blocks with the masked attention map described above. The inference steps are illustrated in Supplementary Figure S1B. During the inference for cell-prompt generation, scGPT generates all genome-wide gene expression conditioned on the specific cell types. A trained cell embedding is inputted at the first position representing the cell type condition. The whole generation process of thousands of gene expression values is conducted in *K* iterative steps (i.e., *K* = 3 steps in Supplementary Figure S1B). For example, in one iteration *i ∈ {*1, 2, … *K}*, the attention masking mechanism allows attention with all predicted genes from previous 0 to *i −* 1 iterations. In each iteration, scGPT selects the top 1*/K* genes from the unknown set with the highest prediction confidence to be included as known genes in the next iteration *i* + 1. Intuitively, this workflow streamlines the generation of gene expression in an auto-regressive manner, where gene expression values with highest prediction confidence are first generated and used to help subsequent rounds of generation. The gene-prompt generation works similarly in an iterative manner. The difference is that it starts with a set of known genes with observed expression values, instead of a cell embedding.

The scGPT attention masking unifies the encoding process of known genes and the generation of unknown genes. It also stands as one of the first transformer schemes to conduct auto-regressive generation for non-sequential data.

#### 4.3.3 Learning objective

We used a **gene expression prediction** objective to optimize the model to predict the expression values for unknown genes. Specifically, we employ a Multi-Layer Perceptron Network (MLP) to estimate the unknown expression values.

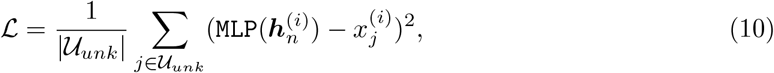

where 𝒰_*unk*_ denotes the set of the output positions for unknown genes, and the 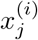 is the actual gene expression value to be predicted. The | *·* | operation retrieves the number of elements of the set.

As mentioned in Online Methods 4.3.2, both gene-prompt and cell-prompt generations are supported. During training, these two modes are conducted consecutively. Among the input gene tokens of one given cell, a proportion of the genes are selected to be the “unknown” genes and their expression values are omitted. Firstly, in the gene-prompt step, the input to the model contains the *< cls >* token embedding, the known gene embeddings, and the unknown gene embeddings. The loss (Equation 10) is computed using the model’s output. Secondly, in the cell-prompt step, the output cell embedding (i.e., 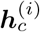 in Online Methods 4.2.2) from the previous step is used to replace the embedding at the *< cls >* position. Other computations remain the same. Finally, the loss values of the two steps are added together and used to compute the gradients to optimize the model parameters.

### 4.4 Fine-tuning objectives

scGPT leverages various fine-tuning objectives to facilitate the learning of biologically valid representations of cells and genes, as well as for regularization purposes such as batch correction.

#### Gene Expression Prediction (GEP)

To encourage the learning of gene-gene interactions, scGPT incorporates gene expression prediction. This fine-tuning objective works similarly to the objective in pre-training (Online Methods 4.3.3), but applies to masked positions instead. To be specific, for each input cell, a subset of gene tokens and their corresponding expression values ***x***^(*i*)^ are randomly masked. scGPT is optimized to accurately predict the expression values at the masked positions. This fine-tuning objective benefits the model in effectively encoding coexpression among the genes in the dataset. The objective minimizes the mean squared error at the masked positions, denoted as ℳ_*mask*_. The GEP works as follows,

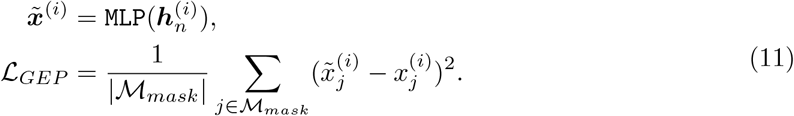

Here, 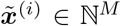 represents the row of expression estimates for cell *i*. Notably, if sequencing batches or modality conditions are provided, we use 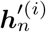 from Equation 8 instead of 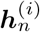.

GEP presents a general self-supervised fine-tuning objective, which aims to forecast gene expression values. In certain downstream tasks, such as perturbation prediction, the model is required to predict perturbed gene expression values instead of the original values. We refer to this variation as **perturb-GEP**. We maintain the MLP estimator in equation 11, but utilize the post-perturbation gene expression as the target 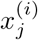. In perturb-GEP, the model is supposed to predict the post-perturbation expression of all input genes.

#### Gene Expression Prediction for Cell Modelling (GEPC)

This fine-tuning objective operates similarly to GEP, but predicts gene expression values based on the cell representation 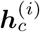 to explicitly facilitate cell representation learning. For each gene *j* in an input cell *i*, we create a query vector ***q***_*j*_ and utilize the parameterized inner product of ***q***_*j*_ and the cell representation 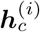 as the predicted expression value.

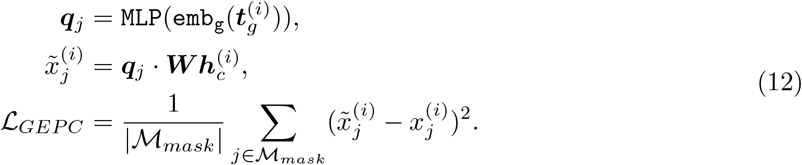

GEPC inherits the gene token embedding, emb_g_ (***t***^(*i*)^*g*), from equation 5. In integration tasks, we utilize 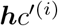 from Equation 9 instead of 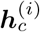. In our experiments, we observed that combining GEP and GEPC leads to significantly improved performance compared to using either method individually.

#### Elastic Cell Similarity (ECS)

This fine-tuning objective enhances cell representations through the utilization of a similarity learning loss [73]:

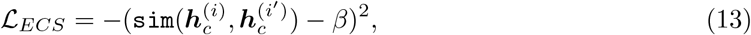

where `sim` represents the cosine similarity function, while *i* and *i*^*′*^ refer to two cells within the minibatch. Additionally, *β* denotes a predefined threshold. The underlying idea behind this approach is to enhance the similarity between pairs exhibiting cosine similarity values above *β*, thereby making them even more similar. Conversely, dissimilar pairs are encouraged to be further apart.

#### Domain Adaptation via Reverse Back-Propagation (DAR)

Cell representation learning is hindered by the presence of batch effects, which result from non-biological batch differences introduced by sequencing technologies [74, 75]. To mitigate this problem, we employ a distinct multi-layer perceptron (MLP) classifier to predict the sequencing batch associated with each input cell, and modify the back-propagation process by reversing the gradients within the model. This approach leverages insights from the robust domain adaptation method proposed by Ganin and Lempitsky [76].

#### Cell Type Classification (CLS)

This fine-tuning objective is designed to leverage the learned cell representations to annotate single cells. We use a separate MLP classifier to predict the cell types from their cell representations 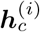. This fine-tuning objective is optimized with the crossentropy loss *ce* between the predicted cell type probabilities and ground-truth labels.

### 4.5 Fine-tuning on downstream tasks

#### Cell type annotation

For the cell type annotation task, we fine-tuned the model on a reference set with ground-truth labels, and validated the annotation performance on a held-out query set. The common set of gene tokens between the pre-trained foundation model and the reference set was retained. The gene expression values were normalized, log-transformed, and binned prior to model fine-tuning. All pre-trained model weights were used to initialize the fine-tuned model, except for the output cell-type classifier which was trained randomly initialized. All gene tokens with both zero and non-zero expression values were used in training. The CLS fine-tuning objective was used to minimize the classification loss.

#### Perturbation response prediction

To fine-tune for the perturbation prediction task, we selected highly variable genes and pre-processed the expression values prior to model training. The parameters of the embedding layers and transformer layers from the pre-trained model were utilized to initialize the fine-tuned model. During fine-tuning, all gene tokens with both zero and non-zero expression values were included. Two notable changes were adopted for the input in the perturbation prediction task: First, we used *log*1*p* transformed expression values as input and target values instead of binned values, to better predict the absolute post-perturbation expression for this task. Second, we appended a binary condition token at each input gene position to indicate whether the gene is perturbed. We adopted the perturb-GEP fine-tuning objective with further modifications to the training setup. Instead of utilizing the masked and unmasked expression values of the same cell as the input and learning target, we employed a control cell as the input and the perturbed cell as the target. This was achieved by randomly pairing a non-perturbed control cell with each perturbed cell to construct the input-target pairs. The input values consisted of all the non-perturbed gene expression values in the control cells. Consequently, the model learned to predict the post-perturbation responses based on the control gene expression.

#### Batch correction on integrating multiple scRNA-seq datasets

Batch effects can be a major confounder in cell type clustering when the input contains multiple datasets from different sequencing batches or technologies. Therefore, we aim to correct batch effects while preserving biological variances when integrating multiple scRNA-seq datasets. For fine-tuning on this integration task, the common set of gene tokens between the pre-trained foundation model and the current dataset was retained. We further selected a subset of highly variable genes from the common set as input. We pre-processed the expression values prior to model training similar to the cell type annotation task. All pre-trained model weights were used to initialize the fine-tuned model. All gene tokens with both zero and non-zero expression values were used in training by default. In addition to GEP and GEPC, the ECS, DAR, and DSBN fine-tuning objectives were optimized simultaneously for enhanced cell contrastive learning and explicit batch correction through reverse back-propagation and domain-specific normalization.

#### Integrative representation learning for scMultiomic data

scMultiomic data may contain different sequencing modalities across experiment batches. We examined two data integration settings, paired and mosaic, for scMultiomic data. In the paired setting, all samples (cells) share all the data modalities sequenced. In the mosaic setting, some batches share a few common data modalities but not all. Due to the presence of additional ATAC and/or protein tokens, we inherited the trained gene embeddings for RNA data only, and trained the additional token embeddings and rest of the model from scratch. Only tokens with non-zero expression values were used in training if the dataset contained additional protein data. Otherwise, both zero and non-zero expression values were used by default. We used an additional set of modality tokens to indicate the data type of each token (i.e., gene, region, or protein) and to facilitate the masked gene and value prediction in GEP and GEPC fine-tuning objectives (Online Methods 4.2.3). The model was optimized with GEP and GEPC fine-tuning objectives by default. If multiple batches are present, DAR was included to facilitate multi-modal batch correction.

#### Gene Regulatory Network Inference

For the gene-embedding-based GRN inference, in the zero-shot setting, we constructed the gene similarity network from the pretrained scGPT’s gene embeddings based on k-nearest neighbors. In the fine-tuned setting, we constructed the gene similarity network in a similar manner from the scGPT model fine-tuned on the Immune Human dataset. Following Ceglia et al. [77]’s pipelines, we further perform Leiden [40] clustering on the similarity graph and extracted gene programs from gene clusters that consist of five or more genes.

For the attention-based target gene selection, we fine-tuned the scGPT blood model on the Adamson perturbation dataset that consists of 87 CRISPR interference experiments on a leukemia cell line [36]. We illustrate the target gene selection pipeline in Figure 6A. For each perturbed gene of interest, we first retrieved two sets of attention maps, perturbed and control, by feeding the model perturbed versus control cell sets respectively. Note that the raw attention scores are obtained from all eight attention heads from the last attention layer of the model. The raw attention scores then go through two rounds of rank normalization, first by row and then by column. The rank-normalized attention scores are then averaged across eight attention heads to output an aggregate attention map. This arrives at the final attention map used for the most influenced gene selection. For each perturbed gene of interest, we select its most influenced genes by ranking the scores from the final attention map in the column of the perturbed gene. This reflects the intuition that the columns in the attention map indicate how much the gene of interest affects the other genes. We offer three most influenced gene selection settings, namely *Control* from the control attention map, *Perturbed* from the perturbed attention map, and *Difference* from the difference between the two. The gene targets selected from the control attention map should reflect the basal pathways that the gene of interest takes part in, whereas the perturbed attention map reflects the post-perturbation effects. The difference between these two attention maps should highlight the most changed edges in the gene network from before to after perturbation. See Online Methods 4.7 for more details on gene network analysis and validation.

### 4.6 Datasets

#### CELLxGENE scRNA-seq human PBMC Collection

We collected data for the whole-human foundation model pre-training from the CELLxGENE portal [28] using the Census API ^2^. We included sequencing protocols of scRNA-seq and snRNA-seq and filtered in samples without disease conditions. This resulted in sequencing data of 33 million cells. For the pre-training of the scGPT blood model specifically, we retrieved over 10.3 million human blood and bone marrow scRNA-seq samples [28]. A total of 65 datasets were collected from CELLxGENE by filtering on Organism (i.e., Homo sapiens), Tissue (i.e., blood, bone marrow), and Disease (i.e., normal, COVID-19, influenza). Additionally, we collected 5.7 million cells of various cancer types to train the pan-cancer model.

#### Multiple Sclerosis

The multiple sclerosis (M.S.) dataset, originally published by Schirmer et al. [78], is accessed from EMBL-EBI (https://www.ebi.ac.uk/gxa/sc/experiments/E-HCAD-35). 9 healthy control samples and 12 M.S. samples are included in the dataset. We split the control samples into the reference set for model fine-tuning and held out the M.S. samples as the query set for evaluation. This setting serves as an example of out-of-distribution data. We excluded three cell types, B cell, T cell, and oligodendrocyte B, that only existed in the query dataset. The final cell counts are 7,844 in the training reference set and 13,468 in the query set. The provided cell-type labels from the original publication were used as ground truth labels for evaluation. The data processing protocol involved selecting highly variable genes (HVG) to retain 3,000 genes.

#### Myeloid

The myeloid (Mye.) dataset [33] can be accessed from the Gene Expression Omnibus (GEO) database using the accession number GSE154763. The dataset consists of nine different cancer types, but for the purpose of training and evaluating the model, six cancer types were selected in the reference set for training, while three cancer types were used for the query set. The reference set contains myeloid cancer types of UCEC, PAAD, THCA, LYM, cDC2, and KIDNEY, while the query set contains MYE, OV-FTC, and ESCA. The dataset was also randomly subsampled. The final cell counts are 9,748 in the reference set and 3,430 in the query set. 3,000 highly variable genes were selected during the data processing.

#### hPancreas

The hPancreas dataset contains five scRNA-seq datasets of human pancreas cells, re-processed by Chen et al. [34] for the cell type annotation task. The five datasets were split into reference and query sets by data sources. The reference set consists of Braon[79] and Muraro[80] datasets, and the query set consists of Xin[81], Segerstolpe[82], and Lawlor[83] datasets. The reference and query sets both have 3,000 genes, and ground-truth annotations retained from their original publications. The reference set contains 10,600 cells of 13 cell groups (alpha, beta, ductal, acinar, delta, PSC, PP, endothelial, macrophage, mast, epsilon, schwann, and t cell). The query set contains 4,218 cells of 11 cell groups (alpha, beta, ductal, PP, acinar, delta, PSC, endothelial, epsilon, mast, MHC class II).

#### PBMC 10K

The PBMC 10K dataset comprises two scRNA-seq batches of human peripheral blood mononuclear cells (PBMCs) obtained from a healthy donor. The dataset was re-processed by Gayoso et al. [44], resulting in 3,346 differentially expressed genes. The first batch contains 7,982 cells, while the second batch contains 4,008 cells. The cell groups annotated using Seurat [42] consist of 9 categories, namely B cells, CD4 T cells, CD8 T cells, CD14+ Monocytes, Dendritic Cells, NK cells, FCGR3A+ Monocytes, Megakaryocytes, and Other.

#### Immune Human

The Immune Human dataset encompasses five scRNA-seq datasets: one derived from human bone marrow and four from human peripheral blood. Various sequencing technologies were employed, including 10X Genomics, 10X Genomics v2, 10X Genomics v3, and Smart-seq2. The dataset comprises a total of 33,506 cells and includes 12,303 genes. The ten distinct batches were defined based on the origin of the donors. The harmonized data contain 16 cell groups. We used the data re-processed and the annotations provided by Luecken et al. [53].

#### Perirhinal Cortex

The Perirhinal Cortex dataset includes two distinct samples, drawn from a larger study by Siletti et al. [45], which originally contained 606 high-quality samples encompassing ten diverse brain regions. Each of the two selected batches from the Perirhinal Cortex dataset comprises a substantial number of cells, with the first batch consisting of 8,465 cells and the second batch containing 9,070 cells. An extensive range of 59,357 genes are incorporated within these datasets. We have made use of the annotations of ten unique cell types provided in the original study.

#### COVID-19

The COVID-19 dataset, derived from work by Lotfollahi et al. [13], is organized into 18 distinct batches and offers a diverse representation of cells from lung tissue, PBMC, and bone marrow. Initially encompassing 274,346 cells and 18,474 genes, this dataset has been subsampled to contain a total of 20,000 cells for the purpose of this study. We made use of the annotations provided by the original study.

#### Adamson

The Adamson perturbation dataset contains gene expression data from the K562 leukemia cell line perturbed by Pertub-seq [36]. This dataset includes 87 unique one-gene perturbations by CRISPR interference, each replicated in around 100 cells.

#### Norman

The Norman perturbation dataset contains gene expression data from the K562 leukemia cell line perturbed by Pertub-seq [37]. This dataset has 131 two-gene perturbations and 105 onegene perturbations. Each perturbation is replicated in around 300-700 cells.

#### 10X Multiome PBMC

The 10X Multiome PBMC dataset [47] contains paired single-cell RNA and ATAC data on human PBMC cells sequenced by the 10X Single Cell Multiome protocol. In this dataset, all samples came from the same healthy donor. Each cell contains both gene expression and chromatin accessibility measurements. The processed data by Cao and Gao [14] contains 9,631 cells with read counts from 29,095 genes and 107,194 regions. The annotations include 19 cell groups (CD14 Mono, CD16 Mono, CD4 Naive, CD4 TCM, CD4 TEM, CD8 Naive, CD8 TEM 1, CD8 TEM 2, HSPC, Intermediate B, MAIT, Memory B, NK, Naive B, Plasma, Treg, cDC, gdT, and pDC).

#### BMMC

The BMMC dataset contains paired single-cell RNA and protein abundance measurements on bone marrow mononuclear cells sequenced by the CITE-seq protocol [48]. These cells came from 12 healthy human donors consisting of 12 batches in this dataset. The processed data contains 90,261 cells with measurements from 14,087 genes and 134 surface proteins. The annotations consist of 45 detailed immune cell subtypes.

#### ASAP PBMC

The ASAP PBMC dataset contains four sequencing batches with three data modalities (gene expression, chromatin accessibility, and protein abundance) [49]. The four batches each contain 5,023, 3,666, 3,517, and 4,849 cells respectively. In batches 1 and 2, all samples have 4,768 genes and 216 protein measurements from CITE-seq. In batches 3 and 4, all samples have 17,742 regions and the same 216 protein measurements from ASAP-seq. The annotations by Mimitou et al. [49] contain 4 cell groups (Bcell, Myeloid, NK, and Tcell).

### 4.7 Benchmarking Experiment Setup

#### scRNA-seq cell type annotation

We benchmarked scGPT against two recent transformer-based cell type annotation methods, scBert [35] and TOSICA [34] on the Mye., M.S., and hPancreas datasets. For each dataset, as described in the previous section, we used the reference data partition for model training and validation. The predicted cell type labels on the query set were retrieved for evaluation. We evaluated cell type assignment performance based on four standard classification metrics, *Accuracy, Precision, Recall*, and *MacroF* 1. *Accuracy, Precision*, and *Recall* are calculated globally for overall performance, whereas *MacroF* 1 is averaged per class to increase the weighing of rare cell types. We also reported a normalized confusion matrix with *Precision* by cell type for additional details. See Supplementary Online Methods S.5 for details on metric calculations.

#### scRNA-seq perturbation

We compared scGPT against the recent perturbation prediction method GEARS [38] and CPA [12]. To ensure consistency, we followed the pre-processing steps outlined by Roohani, Huang, and Leskovec [38] in their benchmark. It is worth noting that the steps are different from the settings reported in CPA. To ensure a fair comparison, we trained all models in the same settings as follows and reported the evaluation results. Initially, gene expression values were normalized per cell using the total counts across all genes, and a logarithmic transformation was applied. Subsequently, we selected 5,000 highly variable genes and incorporated any perturbed genes that were not initially considered into the gene set. In the experiments, for one-gene perturbations in both datasets Adamson et al. [36] and Norman et al. [37], the perturbations are split to ensure that test perturbations are not seen in training, i.e., no cells in the training set has undergone any of the test perturbations. For two-gene perturbations in the Norman et al. [37] dataset, the train-test split consists of three scenarios with increasing difficulty: (1) 0/2 unseen genes, (2) 1/2 unseen genes, and (3) 2/2 unseen genes in the training set.

To evaluate the accuracy of perturbation prediction, we employed the Pearson correlation coefficient (*corr*) between the predicted gene expression and the ground-truth expression values. Additionally, we calculated a variant of the Pearson metric based on the amount of change in expression post-perturbation compared to the control, denoted as *corr(*Δ). We reported these Pearson metrics on the top 20 genes of most changed expression (*DE* genes). Thus, we presented two evaluation metrics in total, namely *corr* and *corr(*Δ) for the *DE* conditions, respectively.

For the cluster-based biological validation, we first retrieved a representative gene expression profile for each perturbation condition from the scGPT model. scGPT predicted the representative perturbation response for each perturbation condition from a single vector of sample control gene expression (i.e., of size 1 *×* M genes), obtained by averaging the gene expression of all control cells in the dataset. The Norman et al. [37] dataset contains 105 unique perturbed genes, which yields a total of 5,565 unique perturbation combinations to predict. We projected the high-dimensional predicted perturbation responses onto a two-dimensional UMAP. We first compared the UMAP against the functional groups found in the original publication by Norman et al. [37], where the 236 perturbation experiments were clustered based on ground-truth perturbation responses and annotated by the marker gene expression for their functional roles. We examined the consistency between the scGPT-predicted UMAP projections and the functional grouping found in the original paper. We then analyzed the sub-clusters present in the scGPT-predicted UMAP. With the Leiden clustering resolution of 0.5, 54 sub-clusters were identified in the UMAP of predicted perturbation responses. We annotated each cluster with the most occurring perturbed gene as its dominant gene.

For the reverse perturbation prediction task, we selected 20 genes to construct a perturbation use case from the Norman dataset and to fine-tune and test a new perturbation model. This subset of 20 genes is selected by maximizing the proportion of ground-truth perturbation data for both training and test cases based on scGPT’s train-test split. This selected subspace contains 39 training cases, 3 validation cases, and 7 test cases out of the 210 unique perturbation combinations. The rest are unseen cases without experiment results. The reverse perturbation prediction follows a topK retrieval task setting: we used the predicted responses from all 210 perturbation conditions as the reference data, and the ground-truth responses from the 7 test cases as the query set. The goal is to retrieve the top perturbation conditions that generate the most similar responses as to a query result. For the reference data, instead of having a single representative gene expression profile for each perturbation condition, we obtained the predicted responses from 40 randomly sampled control cells for added diversity. This yields a reference database containing 8,400 predicted post-perturbation gene expression profiles. For each test case of X+Y perturbation, we used the ground-truth gene expression profiles from all cells that underwent X+Y perturbation as the query set. For TopK retrieval, we designed an ensemble voting strategy that involves two rounds of selection. In the first round, each individual query cell selects its top K most similar expression profiles by Euclidean distance from the reference dataset. In the second round, we rank the candidate perturbation conditions by the number of votes received from all query cells. We report the top K most voted perturbation conditions as the predicted source perturbation conditions from the second round of ranking after ensemble voting. We evaluated the retrieval performance by a modified topK accuracy metric for correct retrieval (i.e., exact match) and relevant retrieval (i.e., matching at least one gene in the actual perturbation combination) as detailed in Supplementary Online Methods S.5.

#### scRNA-seq batch integration

In this work, we compared the performance of scGPT with three other methods, namely Seurat [42], Harmony [43], and scVI [41]. The evaluation covers batch correction and cell type clustering on four integration datasets: COVID-19 [13], PBMC 10K [44], and Perirhinal Cortex [45]. Harmony and scVI are highlighted as the top-performing methods in the recent integration benchmark conducted by Luecken et al. [53]. To ensure a fair comparison, all methods were provided with the same number of 1,200 highly variable genes as input. Gene expression values were normalized per cell by considering the total counts across all genes and subsequently log-transformed. The integrated cell embeddings were obtained after the completion of training and were used for evaluation.

The evaluation of the integrated cell embeddings was performed using biological conservation metrics proposed by Luecken et al. [53]. These metrics include the normalized mutual information (*NMI*_*cell*_), adjusted Rand index (*ARI*_*cell*_), and average silhouette width (*ASW*_*cell*_). These scores measure the consistency between the derived cell type clusters and the ground truth labels. For easier comparison, we also computed the average of these metrics, referred to as *AvgBIO*. Additionally, we reported the batch correction metrics proposed by Luecken et al. [53] to assess batch mixing. The batch correction performance was quantified using the inverse of the average silhouette width for batch clustering, denoted as *ASW*_*batch*_, and the graph connectivity measure, denoted as *GraphConn*. We computed *AvgBATCH* as the average of *ASW*_*batch*_ and *GraphConn* to summarize the batch mixing performance. Furthermore, we introduced an *Overall* score, which is a weighted sum of *AvgBIO* and *AvgBATCH*, consistent with the approach taken by Luecken et al. [53]. See Supplementary Online Methods S.5 for details of metric calculations.

#### scMultiomic integration

We benchmarked scGPT in two integration settings, paired and mosaic, against the recent scMultiomic integration methods Seurat v4 [46], scGLUE [14] and scMoMat [15] respectively. In the paired data integration experiment, we benchmarked scGPT with scGLUE [14] and Seurat v4 [46] on the 10X Multiome PBMC [47] dataset with RNA and ATAC-seq data as the first example. The same 1,200 highly variable genes and 4,000 highly variable peaks were used as input to all methods. We further benchmarked scGPT against Seurat v4 on the BMMC[48] dataset with paired RNA and Protein reads. We did not benchmark scGLUE in this case for fair comparison since the method was not specifically designed to model protein data. Similarly, the same 1,200 highly variable genes and all 134 proteins were used as input. In the mosaic data integration experiment, we benchmarked scGPT with scMoMat [15] on the ASAP PBMC [49] dataset. 1,200 highly variable genes, 4,000 highly variable peaks, and all 216 protein features were used as input to both methods. While keeping the input feature set consistent, we used each method’s custom pre-processing pipeline to normalize the expression values. The integrated cell embeddings were retrieved for evaluation after training.

In all three datasets for both paired and mosaic data integration settings, we evaluated cell embedding quality on the four biological conservation metrics *NMI*_*cell*_, *ARI*_*cell*_, *ASW*_*cell*_, and *AvgBIO*. Since two of the three datasets, BMMC (paired) and ASAP PBMC (mosaic), contain multiple batches, we further evaluated mixing of different omic batches with the three batch correction metrics *ASW*_*batch*_, *GraphConn*, and *AvgBATCH*. An overall score was also reported on the mosaic integration experiment. See Supplementary Online Methods S.5 for details on metric calculation.

#### Gene Regulatory Network Inference

We validated scGPT’s gene embedding similarity network against the known HLA and CD gene networks. For each network, we first defined the related gene set by filtering on gene names with set prefixes (i.e., HLA- and CD-). We then filtered on genes involved in the Immune System R-HSA-168256 pathway from the Reactome 2022 database [54]. For the CD genes, we used the common gene set with the HVGs from the Immune Human dataset for ease of comparison between pre-trained and fine-tuned models. We then extracted gene embeddings of these select genes from the scGPT model and constructed a kNN similarity network. We highlighted sub-networks of strong connections by selecting edges with cosine similarities greater than a certain threshold (i.e., 0.5 for HLA and 0.4 for CD gene network). We then compared the sub-networks against known functional groups from the immune system.

Furthermore, we validated the quality of gene programs extracted by the scGPT model through pathway enrichment analysis. We used each gene program as an input gene list, and selected statistically significant pathways as “pathway hits”. The p-value threshold is adjusted to 0.05 with Bonferroni correction [55] based on the total number of tests performed, i.e., the number of gene programs times the number of pathway tests. We reported the number of pathway hits in the Reactome 2022 database [54]. As a benchmark, we compared the result with the gene programs extracted from the baseline co-expression graph. The co-expression graph was defined by Pearson correlation among genes from their normalized gene expression in the Immune Human dataset. To ensure similar modularity as the scGPT network, we sparsified this graph to a kNN similarity network (k=15). Following the same pipeline as scGPT, we identified gene programs from gene clusters via Leiden clustering. As a sensitivity analysis, we reported the pathway hits for scGPT and the co-expression method at varying Leiden resolutions of 1, 5, 10, 20, 30, 40, 50, and 60. We further examined the common and unique pathways identified by each method at Leiden resolution of 40 to gain more insights into the performance difference.

We validated scGPT’s attention-based most influenced gene selection method in the ChIP-Atlas database [56] which contains experiment-validated gene targets for known transcription factors. We first selected two example transcription factors, *DDIT3* and *BHLHE40*, by cross-checking the perturbed gene list from the Adamson perturbation dataset [36] with ChIP-Atlas. For each transcription factor, we validated the top 20 most influenced genes selected by attention in the *Difference* setting by comparing them against the validated gene targets. Note that in the *Difference* setting, the top 20 genes are selected based on post-perturbation changes by examining the difference between the perturbed attention map and control attention map. The ground-truth gene target list was obtained from ChIP-Atlas by filtering on human genes (hg38) whose transcription start sites lie within 10k-bp distance of the peak-call intervals of the transcription factor. We reported the number of overlaps in the top 20 attention-selected gene targets with the ground-truth target genes.

We subsequently compared the three most influenced gene selection methods (i.e., *Control, Perturbed*, and *Difference*) by examining the overlap between the top 100 genes selected. The overlaps and differences of these three top 100 gene sets are visualized in a Venn diagram. We further validated the pathways these top genes participate in along with the transcription factor in the Reactome database. The pathway hits and the percentage of gene overlap are visualized in a heatmap.

### 4.8 Implementation Details

The pre-trained foundation model has an embedding size of 512. It consists of 12 stacked transformer blocks with 8 attention heads each. The fully connected layer has a hidden size of 512. In pre-training of the whole-human model using 33M cells, we randomly split the data and used 99.7% of the data for training and 0.3% for validation. For the pre-training of other models, including the organ-specific models and the pan-cancer model, we randomly split the data and used 97% of the data for training and 3% for validation. Note that in pre-training, only genes with non-zero expression are input to the model. We set a max input length of 1200. For cells with a number of non-zero genes larger than the max input length, 1200 input genes will be randomly sampled at each iteration. We set the ratio of genes to generate to be uniformly sampled from three options of 0.25, 0.50, and 0.75. The model was optimized by the Adam optimizer, using a mini-batch size of 32, at a starting learning rate of 0.0001 and a 0.9 weight decay after each epoch. The model was trained for a total of 6 epochs.

For the tasks of scRNA-seq batch integration, cell type annotation, and perturbation prediction, we utilized the same model layer configuration inherited from the pre-trained model. During the fine-tuning process, we initiated with a learning rate of 0.0001, which decayed to 90% after each epoch. For the integration task, the mask ratio for GEP and GEPC was set to 0.4, while the parameter *β* in ECS was set to 0.6. When combined with other losses, ECS was assigned a weighting of 10. To divide the datasets into training and validation sets, we employed a ratio of 9:1. The model was trained for a fixed duration of 15 epochs, and after each epoch, the GEP loss value was evaluated on the validation set. The reported results correspond to the model with the best validation score.

For the multi-omic integration task, we loaded the gene embeddings from the pre-trained model and used the same embedding size of 512 for any new tokens (i.e., gene, ATAC-peak, or protein). The main model is set to have 4 stacked transformer blocks with 8 attention heads each, and a hidden layer size of 512. All layers are re-initialized except for the pretrained embedding weights. Each dataset is split into train and evaluation sets at 9:1 ratio. We used DAR weighing of 1.0 for batch integration. We used a starting learning rate of 0.001 and a weight decay of 0.95 after each epoch. We trained the model for fixed 25 epochs with batch size 16 and similarly reported the best-validated model.

We used the `SCANPY` python library [84] for gene expression pre-processing, including normalization, log-transformation, and highly variable gene selection. We used the EpiScanpy python library [85] on chromatin accessibility data for highly variable peak selection. In the scRNA-seq batch integration and scMultiomic integration tasks, the evaluation metrics are calculated using the implementation in `scib.metrics` by Luecken et al. [53]. In the cell-annotation task, the evaluation metrics are implemented using the `scikit-learn` package. In the GRN inference task, the similarity graph construction and Leiden clustering were performed using the `SCANPY` library. The pathway enrichment analysis was implemented using the `GSEApy` package [86].

## Acknowledgement

We would like to express our sincere gratitude to, Dr. Nan Duan, for his invaluable guidance and support throughout the project. We appreciate the valuable feedback from Dr. Lin Zhang during the writing of the manuscript.

## Supplementary Notes

### S.1 Context-Specific Pre-training and Its Influence on Integration Performance

We pre-trained scGPT on cell atlases including all human cell types, envisioning that it contributes to specific downstream applications primarily via fine-tuning. Since downstream applications usually focus on a small subset of tissues or cell types, a natural question for us is whether it would better contribute to a specific fine-tuning task via the pretraining on all cells or a context-specific pretraining. To be specific, for the downstream fine-tuning on specific *contexts* of certain cell types, we are interested in comparing the contributions of three different pretrained models: (1) scGPT (whole-human), which is the model pretrained generally on all human cell types; (2) in-context models, which during pretraining have seen similar cell types as in the downstream applications; (3) out-of-context models, which are pretrained largely on different tissues or cell types from the downstream applications. To conduct this comparison, we tested a range of pre-trained models on the scRNA-seq data integration task. These pre-trained models were originally trained using distinct tissue-specific datasets, including lung, blood, heart, kidney, brain, pancreas, intestine, and an all-inclusive, whole-human cells dataset. In particular, we subsampled from the original whole-human dataset to generate a dataset of 13.2 million cells, aligning it in size with the blood (10.3 million cells) and brain (13.2 million cells) datasets for a more direct comparison of context impact. We employed each of the eight pre-trained models to perform data integration on the COVID-19 dataset [13], followed by a comparative performance analysis. Given its diverse cellular composition, including Lung, PBMC, and Bone Marrow cells, this dataset provides an ideal platform for investigating the effects of employing pre-trained models from different cellular contexts.

Our process yielded notable results, revealing a clear correlation between the relevance of the model context used in the pre-training initiatives and their subsequent performance on the COVID-19 dataset. Supplementary Figure S8 graphically illustrates these findings, showcasing the average *AvgBIO* score along with the standard error derived from five integration experiments for each pretrained model (Panel A). Moreover, the UMAP visualization (Panel B) presents an in-depth view of the cell embeddings colored by cell types, substantiating the quality of the learned representations and visually validating the models’ integration performance. Notably, the top performers in this analysis were models pre-trained on whole-human, blood, and lung datasets, which correspond closely to the cell types present in the COVID-19 dataset.

In particular, even though the brain pre-trained model was trained on a substantial dataset of 13 million cells, it trailed in performance by 8% compared to the blood pre-trained model with a similar dataset size. This gap in performance sheds light on the importance of cellular context relevance. Specifically, the cellular context of the blood model aligns more closely with the COVID-19 dataset, which includes immune cells, bone marrow cells, lung cells, and PBMCs. Thus, it becomes evident that the alignment of the cellular context in the pre-training phase plays a critical role in achieving superior results for downstream data integration tasks, even when datasets of similar sizes are compared. In light of our findings, the whole-human pre-trained model, embodying a vast spectrum of cell types, consistently demonstrates robust performance across diverse analyses.

Our investigation underlined the significance of cellular context in single-cell RNA-seq data integration tasks. In certain circumstances, when the cellular context of the target dataset aligns with the tissue-specific pre-trained models, these models can excel. Overall, while it is essential to consider the cellular context, the whole-human pre-trained model emerges as a versatile and reliable option for a wide range of applications.

### S.2 Benchmarking results on downstream tasks

**Table S1:** Cell Type Annotation Benchmark Results. scGPT was benchmarked with TOSICA [34] and scBert [35] on the Myeloid (Mye.), Multiple Sclerosis (M.S.), and hPancreas [34] datasets for cell type annotation performance. We present four classification evaluation metrics *Accuracy, Precision, Recall*, and *MacroF* 1. See metric details in Supplementary Online Methods S.5.

| Dataset | Model | Classification Metrics |  |  |  |
| --- | --- | --- | --- | --- | --- |
|  |  | *Accuracy* | *Precision* | *Recall* | *MacroF1* |
| Myeloid | scGPT (fine-tuned) | **0.642** | **0.366** | **0.347** | **0.346** |
|  | scGPT (from-scratch) | 0.606 | 0.304 | 0.339 | 0.309 |
|  | TOSICA | 0.488 | 0.316 | 0.276 | 0.275 |
|  | scBert | 0.525 | 0.331 | 0.323 | 0.298 |
| Multiple Sclerosis | scGPT (fine-tuned) | **0.856** | **0.729** | **0.720** | **0.703** |
|  | scGPT (from-scratch) | 0.798 | 0.660 | 0.623 | 0.600 |
|  | scBert | 0.785 | 0.604 | 0.624 | 0.599 |
|  | TOSICA | 0.758 | 0.664 | 0.585 | 0.578 |
| hPancreas | scGPT (fine-tuned) | **0.968** | **0.735** | **0.725** | **0.718** |
|  | scGPT (from-scratch) | 0.936 | 0.665 | 0.668 | 0.622 |
|  | TOSICA | 0.960 | 0.661 | 0.681 | 0.656 |
|  | scBert | 0.964 | 0.699 | 0.689 | 0.685 |

**Table S2:** scRNA-seq Integration Benchmark Results. scGPT was benchmarked with scVI [41], Seurat [42], and Harmony [43] on the COVID-19 (18 batches) [13], PBMC 10K (2 batches) [44] and Perirhinal Cortex (2 batches) [45] datasets for cell type clustering and batch correction performance. We present three aggregate scores *AvgBIO, AvgBATCH*, and *Overall*. These aggregate scores were calculated from three detailed biological conservation metrics (*NMI*_*cell*_, *ARI*_*cell*_, *ASW*_*cell*_) and two batch correction metrics (*ASW*_*batch*_, *GraphConn*). See metric details in Supplementary Online Methods S.5

| Dataset | Model | Biological Conservation |  |  |  | Batch Correction |  |  | Overall |
| --- | --- | --- | --- | --- | --- | --- | --- | --- | --- |
| | | AvgBIO | $NMI_{cell}$ | $ARI_{cell}$ | $ASW_{cell}$ | AvgBATCH | $ASW_{batch}$ | $GraphConn$ | |
| COVID-19 [13] | scGPT (fine-tuned) | **0.504** | 0.659 | 0.400 | 0.452 | **0.850** | 0.826 | 0.874 | **0.642** |
|  | scGPT (from-scratch) | 0.450 | 0.602 | 0.318 | 0.429 | 0.744 | 0.712 | 0.776 | 0.567 |
|  | scVI [41] | 0.502 | 0.638 | 0.408 | 0.461 | 0.838 | 0.833 | 0.844 | 0.637 |
|  | Seurat [42] | 0.413 | 0.513 | 0.289 | 0.437 | 0.790 | 0.799 | 0.781 | 0.564 |
|  | Harmony [43] | 0.327 | 0.482 | 0.185 | 0.313 | 0.680 | 0.642 | 0.720 | 0.468 |
| PBMC 10K [44] | scGPT (fine-tuned) | **0.821** | 0.850 | 0.873 | 0.740 | 0.923 | 0.950 | 0.895 | **0.862** |
|  | scGPT (from-scratch) | 0.723 | 0.738 | 0.793 | 0.639 | 0.893 | 0.919 | 0.867 | 0.791 |
|  | scVI | 0.753 | 0.819 | 0.847 | 0.592 | **0.947** | 0.967 | 0.928 | 0.831 |
|  | Seurat | 0.724 | 0.808 | 0.722 | 0.641 | 0.940 | 0.960 | 0.920 | 0.810 |
|  | Harmony | 0.784 | 0.860 | 0.902 | 0.591 | 0.940 | 0.975 | 0.906 | 0.846 |
| Perirhinal Cortex [45] | scGPT (fine-tuned) | **0.899** | 0.930 | 0.919 | 0.848 | 0.930 | 0.898 | 0.964 | 0.911 |
|  | scGPT (from-scratch) | 0.889 | 0.886 | 0.895 | 0.886 | 0.884 | 0.892 | 0.875 | 0.887 |
|  | scVI | 0.869 | 0.980 | 0.990 | 0.637 | 0.966 | 0.939 | 0.992 | 0.908 |
|  | Seurat | 0.878 | 0.914 | 0.897 | 0.822 | 0.965 | 0.938 | 0.992 | 0.913 |
|  | Harmony | 0.890 | 0.960 | 0.960 | 0.749 | **0.975** | 0.957 | 0.992 | **0.924** |

**Table S3:**
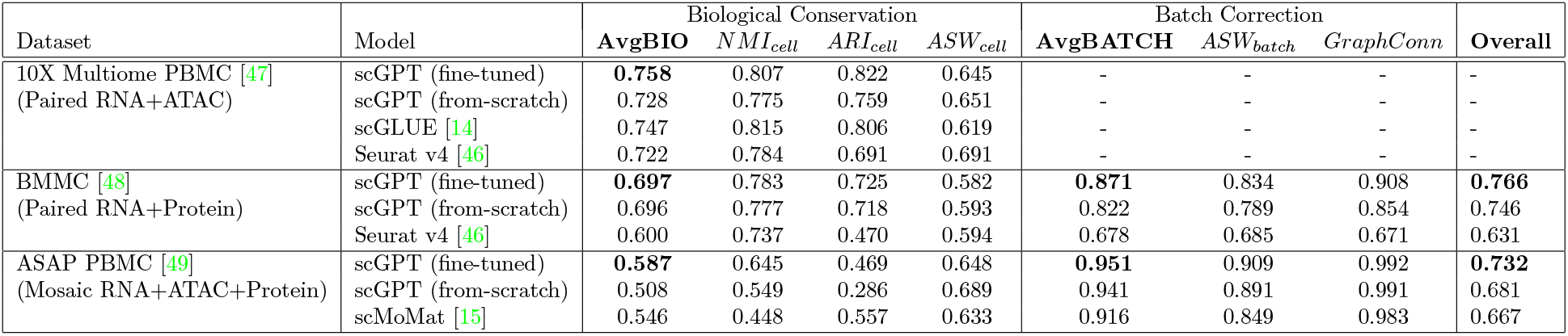
scMultiomic Integration Benchmark Results. For the paired 10X Multiome PBMC[47] dataset, scGPT was benchmarked with scGLUE[14] and Seurat v4[46] for cell type clustering performance evaluated on four biological conservation metrics. The data has only one technical batch. Batch correction metrics are not applicable to this setting. For the paired BMMC[48] and mosaic ASAP PBMC[49] datasets, scGPT was benchmarked with Seurat v4 and scMoMat [15] respectively on cell type clustering and multi-omic integration performance, evaluated on eight biological conservation and batch correction metrics.

| Dataset | Model | Biological Conservation |  |  |  | Batch Correction |  |  | Overall |
| --- | --- | --- | --- | --- | --- | --- | --- | --- | --- |
| | | AvgBIO | $NMI_{cell}$ | $ARI_{cell}$ | $ASW_{cell}$ | AvgBATCH | $ASW_{batch}$ | GraphConn | |
| 10X Multiome PBMC [47]<br>(Paired RNA+ATAC) | scGPT (fine-tuned) | **0.758** | 0.807 | 0.822 | 0.645 | - | - | - | - |
|  | scGPT (from-scratch) | 0.728 | 0.775 | 0.759 | 0.651 | - | - | - | - |
|  | scGLUE [14] | 0.747 | 0.815 | 0.806 | 0.619 | - | - | - | - |
|  | Seurat v4 [46] | 0.722 | 0.784 | 0.691 | 0.691 | - | - | - | - |
| BMMC [48]<br>(Paired RNA+Protein) | scGPT (fine-tuned) | **0.697** | 0.783 | 0.725 | 0.582 | **0.871** | 0.834 | 0.908 | **0.766** |
|  | scGPT (from-scratch) | 0.696 | 0.777 | 0.718 | 0.593 | 0.822 | 0.789 | 0.854 | 0.746 |
|  | Seurat v4 [46] | 0.600 | 0.737 | 0.470 | 0.594 | 0.678 | 0.685 | 0.671 | 0.631 |
| ASAP PBMC [49]<br>(Mosaic RNA+ATAC+Protein) | scGPT (fine-tuned) | **0.587** | 0.645 | 0.469 | 0.648 | **0.951** | 0.909 | 0.992 | **0.732** |
|  | scGPT (from-scratch) | 0.508 | 0.549 | 0.286 | 0.689 | 0.941 | 0.891 | 0.991 | 0.681 |
|  | scMoMat [15] | 0.546 | 0.448 | 0.557 | 0.633 | 0.916 | 0.849 | 0.983 | 0.667 |

**Table S4:**
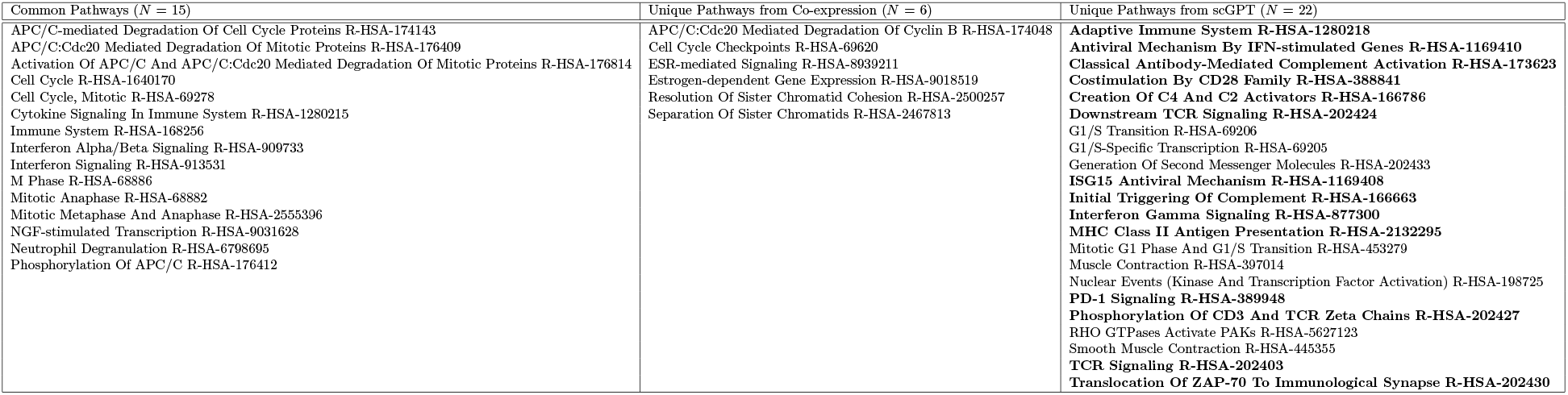
Comparison of Common and Unique Pathways Identified by scGPT and the Coexpression Network From the Reactome Database. The enriched pathways from gene programs extracted by both methods at Leiden resolution 40 are listed here for comparison, corresponding to the Venn diagram in Figure 5 E. The 14 immune-related pathways uniquely identified by scGPT are highlighted in bold.

### S.3 Enhancing Speed and Decreasing Memory Usage in Fine-tunings

In an attempt to hasten the fine-tuning phase and render scGPT more accessible for users, we explored a variety of fine-tuning methods in the context of the scRNA-seq integration task. The pre-trained weights from the whole-human dataset were employed to initialize our entire model. As the standard practice, the baseline Full fine-tune procedure involves gradient updates for all model parameters and includes all zero-expressed genes. We undertook experiments that involved freezing the embedding layers and omitting zero-expressed genes during fine-tuning. From data presented in table S5, we noticed a significant drop in the training time per epoch as well as in GPU Memory usage throughout the fine-tuning process. Specifically, the removal of zero-expressed genes resulted in a substantial reduction in our model’s maximum sequence length to approximately 40 *∼* 60% of its initial length, effectively halving both the time spent on training epochs and peak GPU Memory utilization. Furthermore, by freezing the embedding layer, we achieved an additional reduction in Peak GPU Memory usage by 1GB, a slight increase in training speed, while preserving a comparable AvgBio score.

**Table S5:** Benchmarking Results for scRNA-seq Integration Speed Enhancement Techniques. scGPT was evaluated based on different fine-tuning strategies on four datasets: COVID-19 (18 batches) [13], PBMC 10K (2 batches) [44], and Perirhinal Cortex (2 batches) [45]. These evaluations were performed to assess cell type clustering and batch correction performance. This table encapsulates two system metrics: the average duration of training per epoch and the maximum GPU memory usage on an A100 GPU. The resulting *AvgBio* score is also presented for performance comparisons.

| Dataset | Fine-tuning Option | Fine-tuning Metrics |  | AvgBio |
| --- | --- | --- | --- | --- |
|  |  | Avg Epoch Time(s) | Peak GPU Mem(G) |  |
| COVID-19 [13] | Default | 93.89 | 18.784 | **0.504** |
|  | Accelerated | **28.84** | **7.848** | 0.473 |
| PBMC 10K [44] | Default | 56.91 | 18.248 | 0.821 |
|  | Accelerated | **26.40** | **8.088** | **0.828** |
| Perirhinal Cortex [45] | Default | 81.00 | 18.248 | **0.899** |
|  | Accelerated | **37.22** | **7.816** | **0.899** |

### S.4 Comparison to existing approaches

#### Transformers for modelling single-cell sequencing data

Transformer models employing self-attention mechanisms [27] have demonstrated remarkable success in the field of natural language processing (NLP) [64], computer vision [87], and protein folding prediction [88]. However, there have been few efforts to incorporate the transformer architecture into single-cell biology and its related applications. Shen et al. [89] utilize a transformer decoder setup to learn the sequence of the names of highly expressed genes, but they do not take into account the actual sequenced expression abundance, resulting in the loss of crucial biological information. scBERT [35] and TOSICA [34] used BERT-like architectures [64] to train cell embeddings but only applied the model on the supervised task of cell annotation. Similarly, Connell, Khan, and Keiser [90] utilized transformer encoders mainly to predict genetic perturbation responses. To our best knowledge, scGPT is one of the first methods to provide a generative pre-trained transformer foundation model for multiple single-cell analysis tasks.

#### Pre-training on large cell atlases

Although the idea of employing pre-training and finetuning on a wide range of downstream tasks as a unified framework remains largely unexplored, several works have attempted to use transfer learning on specific tasks. scArches [13] devised a transfer-learning-based approach for reference mapping by pre-training a conditional variational autoencoder (VAE) on the reference datasets. However, the scale of the reference datasets remains limited, and the VAE-based architecture does not incorporate attention computation. On the other hand, scBERT [35] is pre-trained on 1 million cells with a BERT architecture. However, the downstream application focuses on cell type annotation only, thus restricting the generalizability of the pre-training and fine-tuning strategy. A recent work, Geneformer [91], has extended pretrained transformers beyond cell type annotation to gene network analysis, but the work didn’t demonstrate abilities for perturbation response prediction or multi-omic integration. scGPT has compiled an unprecedented scale of pre-training data and evaluated on a diverse range of downstream tasks, which presents the pre-training and fine-tuning strategy as a unified framework for versatile single-cell analysis.

#### Learning cell and gene representation for scRNA-seq on downstream tasks

Cell representation learning facilitates a variety of downstream tasks such as cell type annotation, multi-omic integration, and perturbation prediction. A popular framework Seurat [42, 92] employs nearest-neighbor-based alignment to remove batch effect via linear transformation in the embedding space. LIGER [93] and OCAT [94] use matrix factorization to extract latent cell embeddings. Recently, Deep Learning methods and especially VAE-based generative models have gained increasing popularity, as they generate deep embeddings via non-linear transformations through neural networks [95]. scVI [41] learns latent cell representations by reconstructing original gene expression via variational inference. TotalVI [96], scGen [11], CPA [12] and MultiCPA [97] utilized similar models and extended the application to multi-omics and perturbation prediction. On the other hand, gene representation learning also supports many downstream tasks including gene regulatory network and functional pathway analysis. As an example, GeneVector [77] detects gene-gene functional relations by factorizing the co-expression and mutual information matrix of the sequencing readout. VEGA [98] utilizes a sparse VAE architecture to encode gene network activities for added interpretability. DeepMAPS [99] utilizes graph neural networks to encode cell and gene nodes for related tasks. Despite the importance of the two branches of research in cell and gene embedding learning, few approaches have worked on jointly learning both. scGPT stands out as an approach to effectively learn both embeddings of cells and genes jointly in a shared architecture.

### S.5 Evaluation Metric Calculations

#### S.5.1 Cell Type Assignment

We used the standard classification metrics *Accuracy, Precision, Recall*, and *MacroF* 1 to evaluate cell type assignment performance. The *Accuracy, Precision, Recall*, and *MacroF* 1 scores are calculated from true positives (*tp*), false positives (*fp*), true negatives (*tn*), and false negatives (*fn*) globally or averaged per class.

The *Accuracy, Precision* and *Recall* scores are calculated globally:

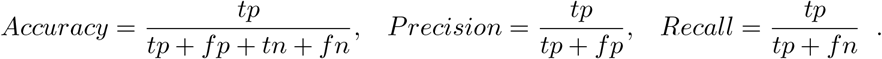

The *MacroF* 1 score is calculated per cell type *c* first and averaged across cell types:

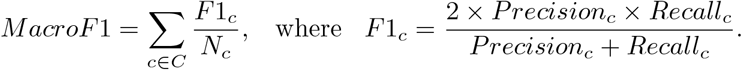

The above metrics are calculated using scikit-learn’s implementations [100].

#### S.5.2 Reverse Perturbation - Predicting driving gene perturbations with TopK Retrieval

We employed two modified topK retrieval accuracy metrics to assess the reverse perturbation prediction performance. The hit rate of correct predictions calculates the proportion of test cases where the topK retrieved experiments contain the target (i.e., query) condition. For example, for each test case of target condition X+Y, if the topK retrieved experiments contain X+Y, we count this test case as a hit. We also reported a relaxed topK accuracy metric for relevant retrievals with one-gene overlap. The percentage of test cases with relevant predictions calculates the proportion of test cases where the topK retrieved experiments contain any cases with a one-gene overlap with the target condition. For example, for the same test case X+Y, if the topK retrieved experiments contain X, Y, X+A, A+X, Y+A, or A+Y, we count this test case as a hit. This relaxed metric aims to provide added interpretability for scGPT’s choices of retrieval.

#### S.5.3 Single-cell integration

We adopted the evaluation metric calculations outlined by Luecken et al. [53] in their benchmark study. Each metric is described below.

##### Normalized Mutual Information

To quantify the concurrence between the cell type labels based on ground truth and the Louvain cluster labels obtained from integrated cell embeddings, we computed the normalized mutual information (NMI) score. The Louvain clustering was conducted across resolutions ranging from 0.1 to 2, with increments of 0.1. The best score will be selected. The NMI score for cell types, referred to as ***NMI*_*cell*_**, ranges between 0 and 1, where a higher score indicates a better match of cell types.

##### Adjusted Rand Index

The adjusted rand index (ARI) was employed to assess both the agreement between the annotated labels and the MNI-optimized Louvain clusters. Furthermore, the rand index was adjusted to account for randomly correct labels. The ARI score for cell types, denoted as ***ARI*_*cell*_**, ranges from 0 to 1, where 0 corresponds to random labeling and 1 represents a perfect match.

##### Average Silhouette Width

The silhouette width assesses the relationship between a cell’s within-cluster distances and its distances to the closest cluster boundaries. By averaging the silhouette widths of all cells, we calculate the average silhouette width (ASW) score. This score ranges from -1 to 1, where a score of 1 indicates well-separated clusters, while scores from -1 to 0 suggest overlapping clusters and misclassification.

For evaluating cell type clustering, we compute the ASW score based on cell type labels, represented as ***ASW*_*cell*_**. To obtain this score, we utilize the following formula:

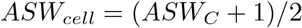

Here, *C* represents the cell types.

Regarding batch mixing evaluation, we calculate the ASW score considering batch labels and adjust it by subtracting 1. This score is denoted as ***ASW*_*batch*_**. The calculation is as follows:

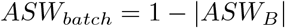

Both ***ASW*_*cell*_** and ***ASW*_*batch*_** have values between 0 and 1. Higher scores indicate better cell-type clustering or batch-mixing performance.

##### Graph Connectivity

The graph connectivity metric quantifies the average proportion of cells within each cell type that are connected through a kNN (k-nearest neighbors) graph. For every cell identity *c* in the set *C*, we compute the size of the largest connected component using kNN among cells exclusively belonging to identity *c*. This value is divided by the total number of cells with identity *c* to obtain a normalized measure. The ***GraphConn*** score is then reported as the average across all cell types:

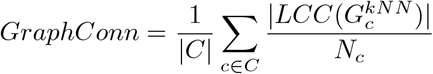

Here, *LCC* represents the largest connected component, and *N* denotes the number of cells of each celltype.

##### Aggregated Metrics

The aggregated metric ***AvgBIO*** calculates the average of biological conservation metrics:

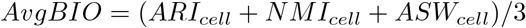

Similarly, the aggregated metric ***AvgBAT CH*** computes the average of batch mixing metrics:

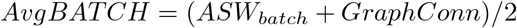

In accordance with the convention established in [53], an ***Overall*** metric is derived as the weighted average of ***AvgBIO*** and ***AvgBAT CH***:

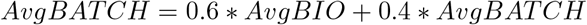

## Supplementary Figures

**Figure S1:**
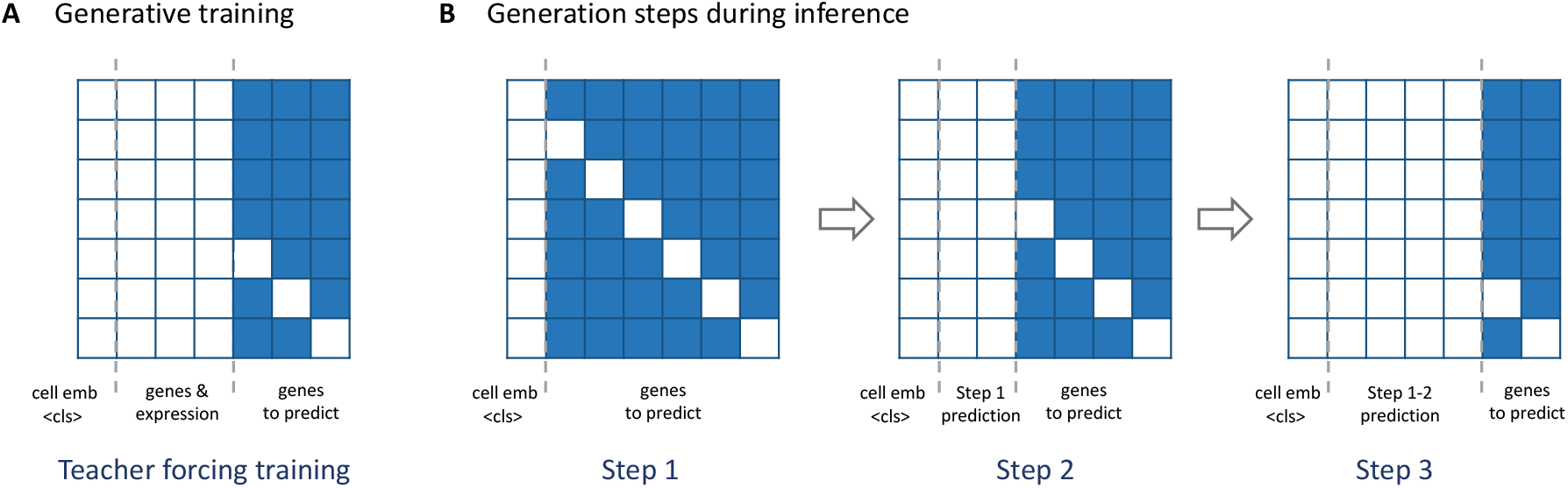
The scGPT Attention Mask. The masked positions are colored in blue, and the unmasked positions in white. These masked and unmasked positions correspond to the *M × M* attention map for *M* input tokens. The row indices correspond to queries and columns correspond to keys. In the self-attention computation of transformers, the attention scores on the masked positions will be removed. The token identity associated with each column is annotated below, namely “cell emb *< cls >*“for cell embedding, “genes & expression” for known genes, and “genes to predict” for unknown genes. *(A)* scGPT attention mask in training where only query gene and the known genes participate in attention computation. *(B)* After training, the attention mask at each step during the iterative process of scGPT cell-prompt generation.

**Figure S2:**
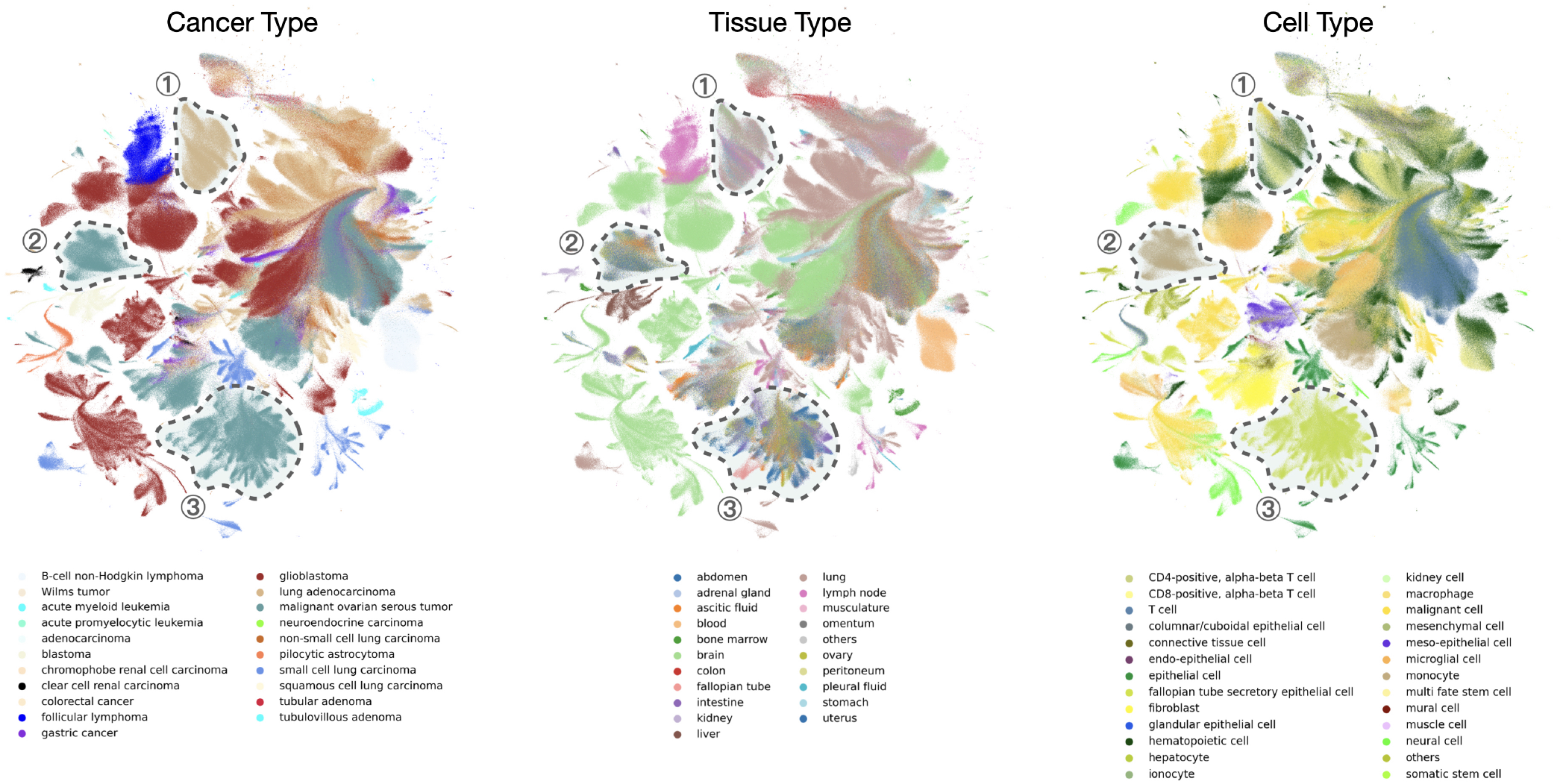
UMAP of 3 million cancer cells using the cell embeddings from the pre-trained pancancer model. From left to right, the colors indicate the cancer types, tissue types, and cell types. We observed that the model is able to generate cell embeddings revealing the difference in cancer and cell types primarily, exemplified by the outlined three regions.

**Figure S3:**
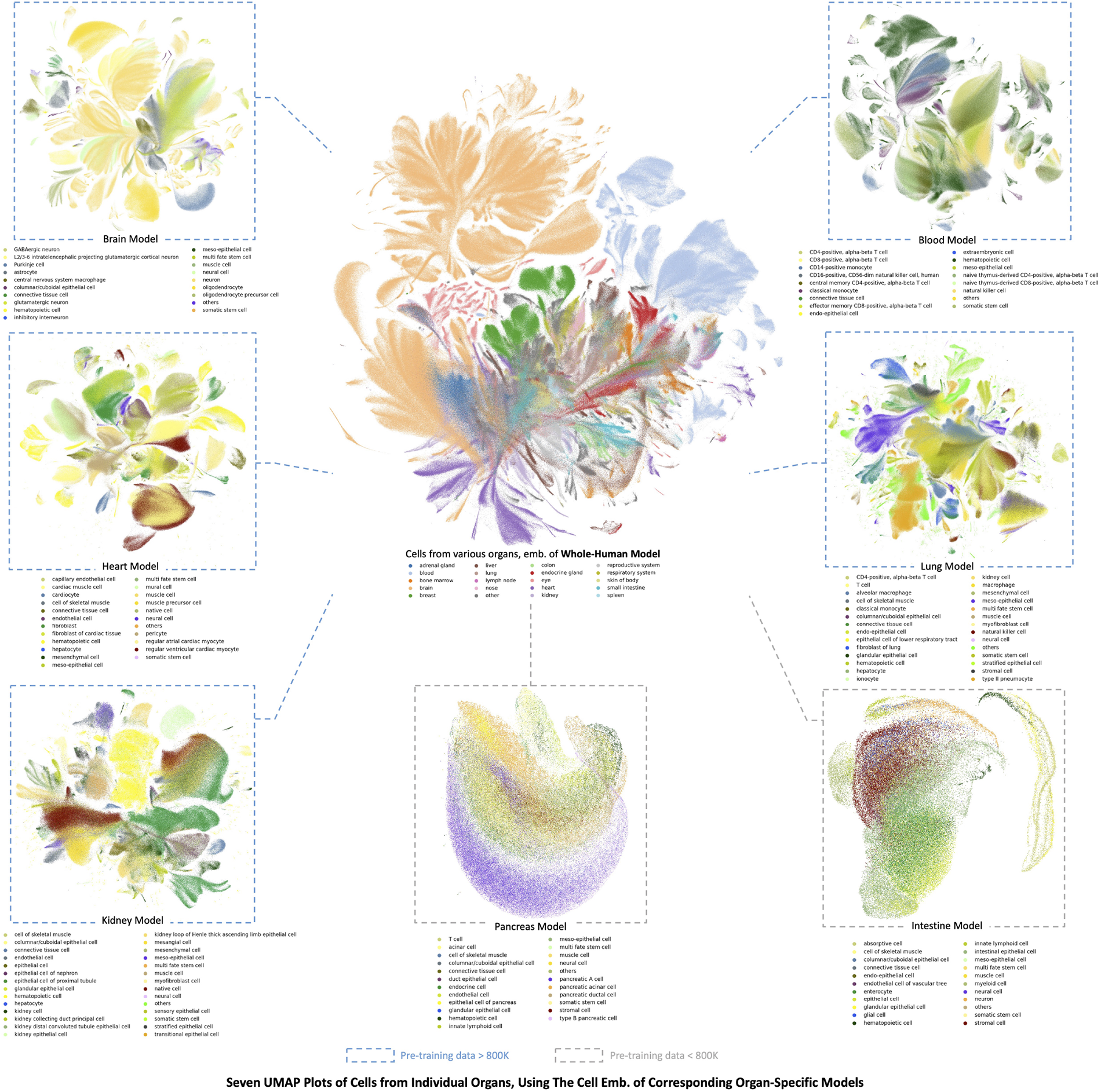
Organ-specific models. (*Center*) The UMAP visualization of selected 3 million collected normal human cells using the cell embeddings from the pre-trained scGPT whole-human model. Cells are colored by the organs of origin. (*Around*) The UMAP visualization of cells from each organ using the cell embeddings from the corresponding organ-specific models. The colors in each image indicate major cell types. For example, the top left UMAP visualizes brain cell embeddings from the scGPT model that was specifically pre-trained on brain cells. The outline color of each UMAP plot indicates whether the size of the organ-specific training data is larger than 800,000 cells (blue) or not (grey). We observed that models trained on sufficient data (i.e., *>* 800,000 cells) could generate decent cell embeddings that can separate major cell types.

**Figure S4:**
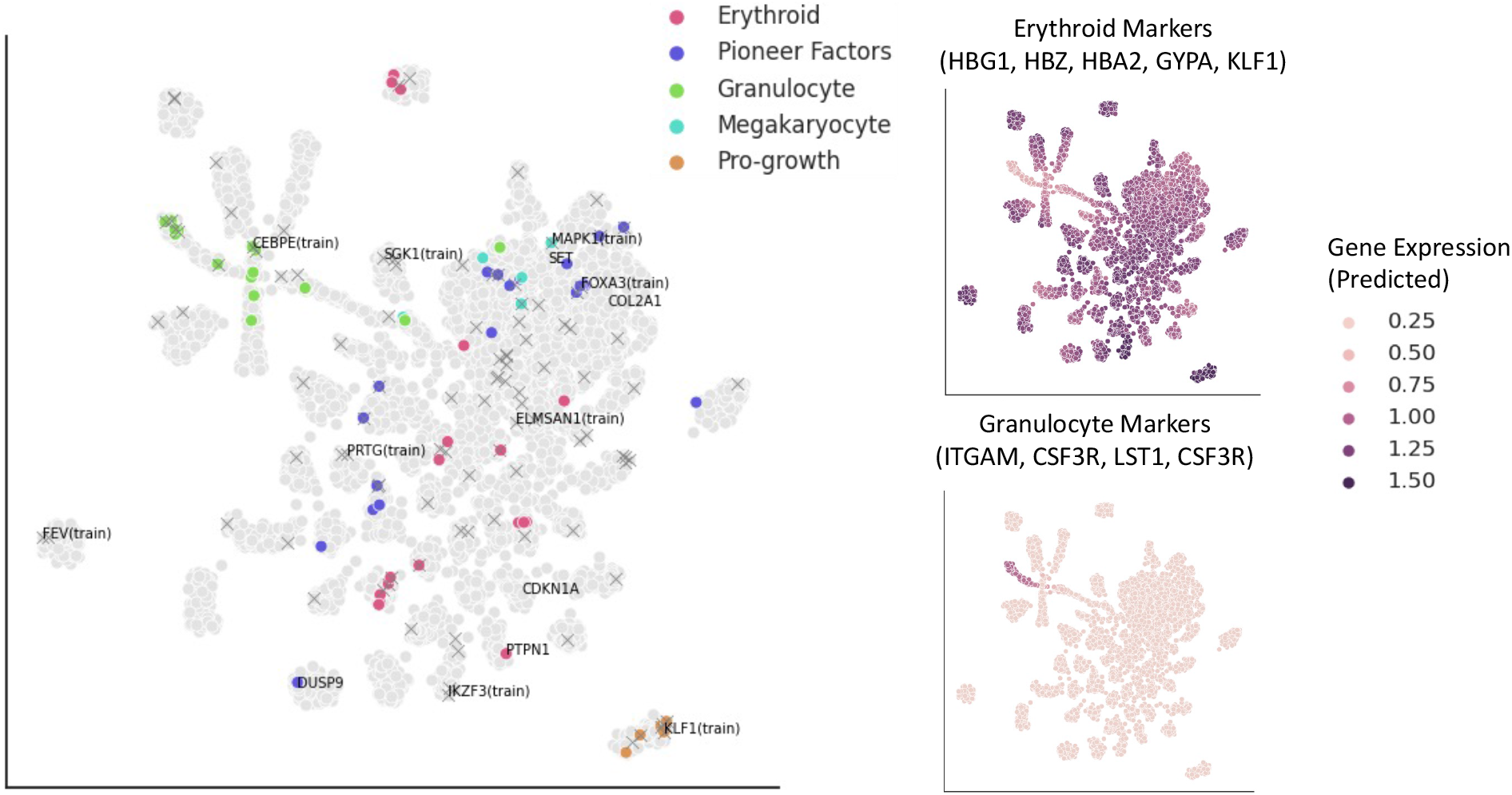
Visualization of Annotated Functional Groups by Norman et al. [37]. *(Left)* UMAP visualization of perturbation condition embeddings colored by *functional groups* on the left. Crosses indicate perturbations that have been tested experimentally in the original study. Colored dots indicate perturbation conditions with annotations. *(Right)* UMAP visualization of perturbation condition embeddings colored by average predicted marker gene expression on the right for the Erythroid and Granulocyte cell groups.

**Figure S5:**
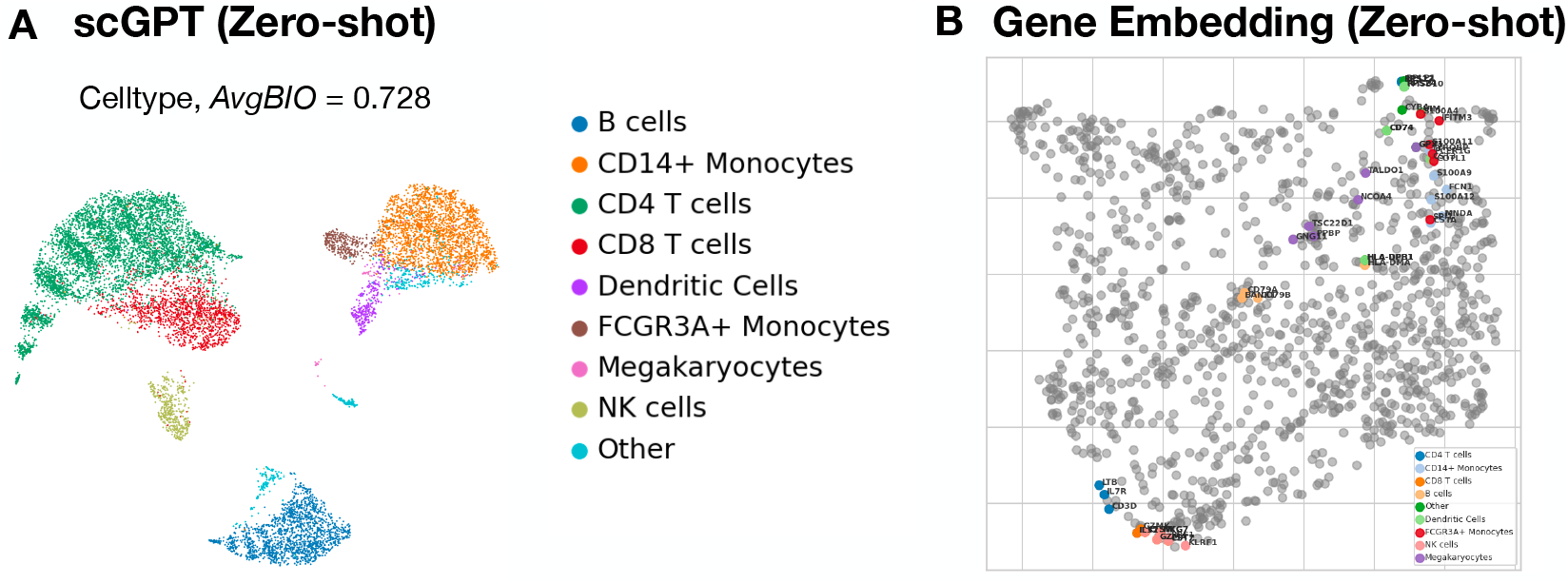
Visualization of the scGPT cell and gene embeddings on the PBMC 10K dataset, using the pre-trained model without fine-tuning (i.e., zero-shot). *(A)* UMAP visualization of cell embeddings colored by *cell types. (B)* UMAP visualization of gene embeddings. The highly variable genes corresponding to major *celltype* were colored accordingly.

**Figure S6:**
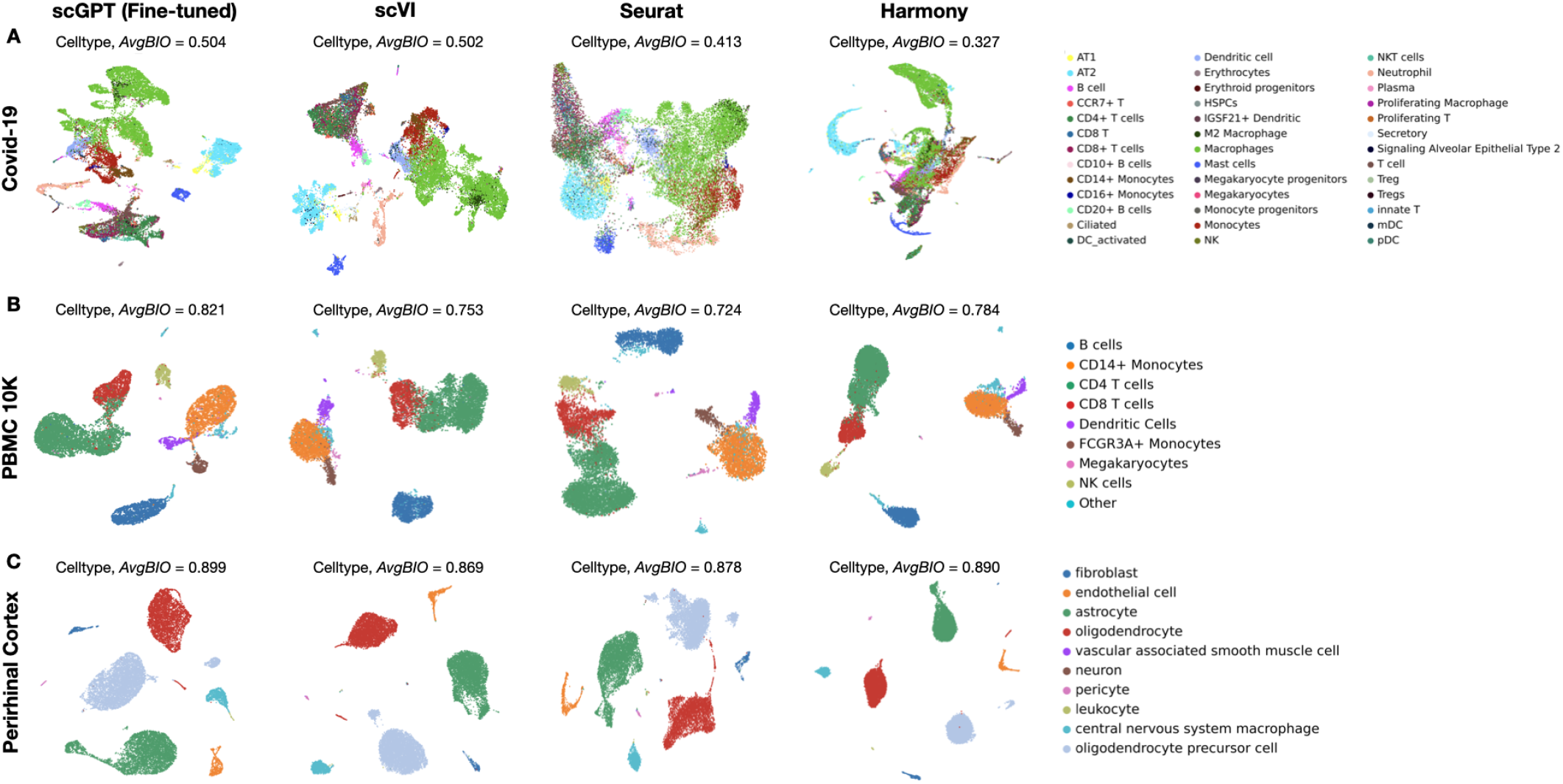
*(A,B,C)* Benchmark of the fine-tuned scGPT model with scVI [41], Seurat Seurat [42], and Harmony Harmony [43] on the COVID-19 (18 batches) [13], PBMC 10K (2 batches) [44], and Perirhinal Cortex (2 batches) [45] datasets for cell type clustering performance upon batch integration. The UMAP plot of learned cell embeddings was colored by *cell types*.

**Figure S7:**
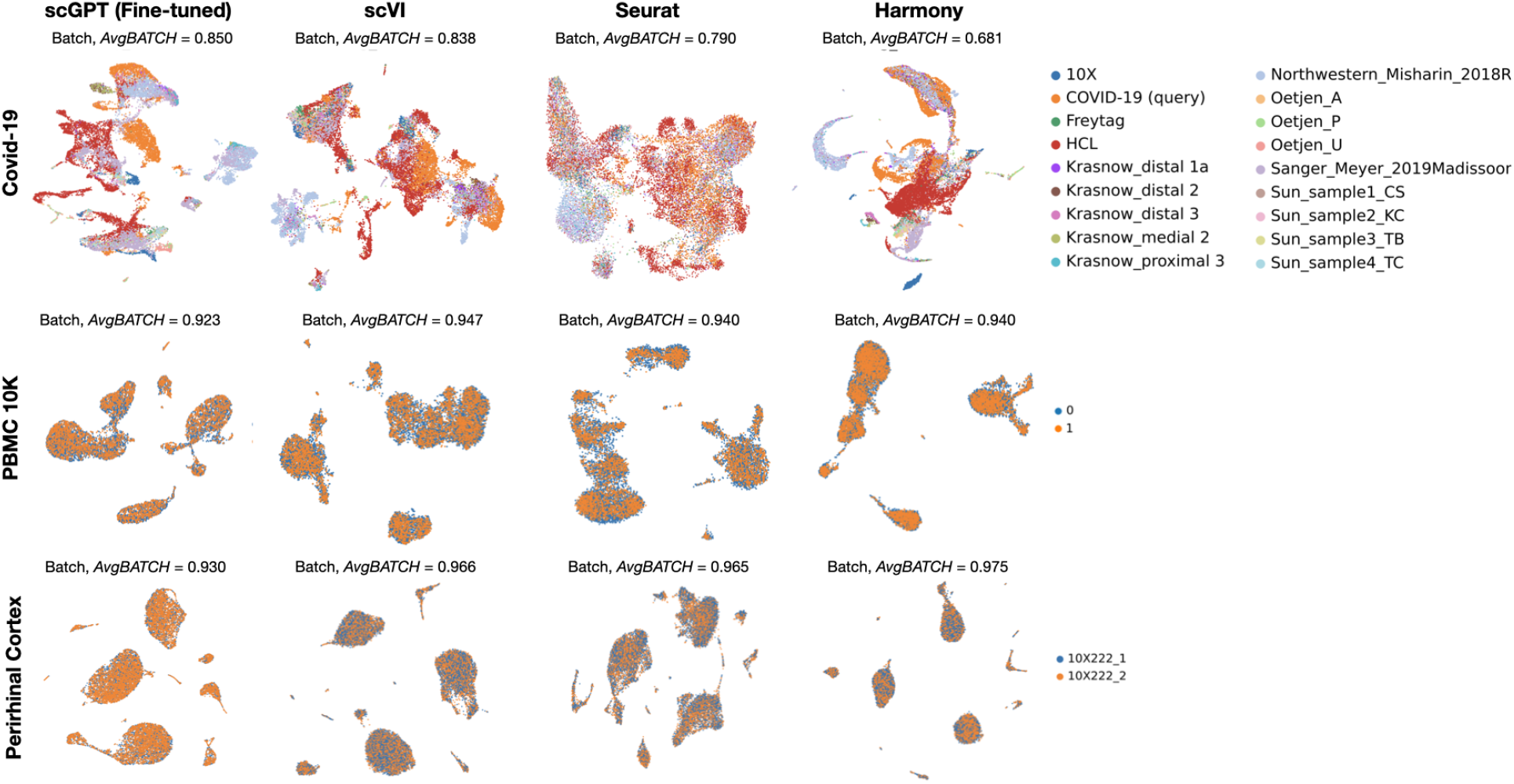
Benchmark of scGPT with scVI [41], Seurat [42], and Harmony [43] on the COVID-19 (18 batches) [13], PBMC 10K (2 batches) [44], and Perirhinal Cortex (2 batches) [45] Datasets for Batch Correction. UMAP visualization of cell embeddings colored by *sequencing batches*.

**Figure S8:**
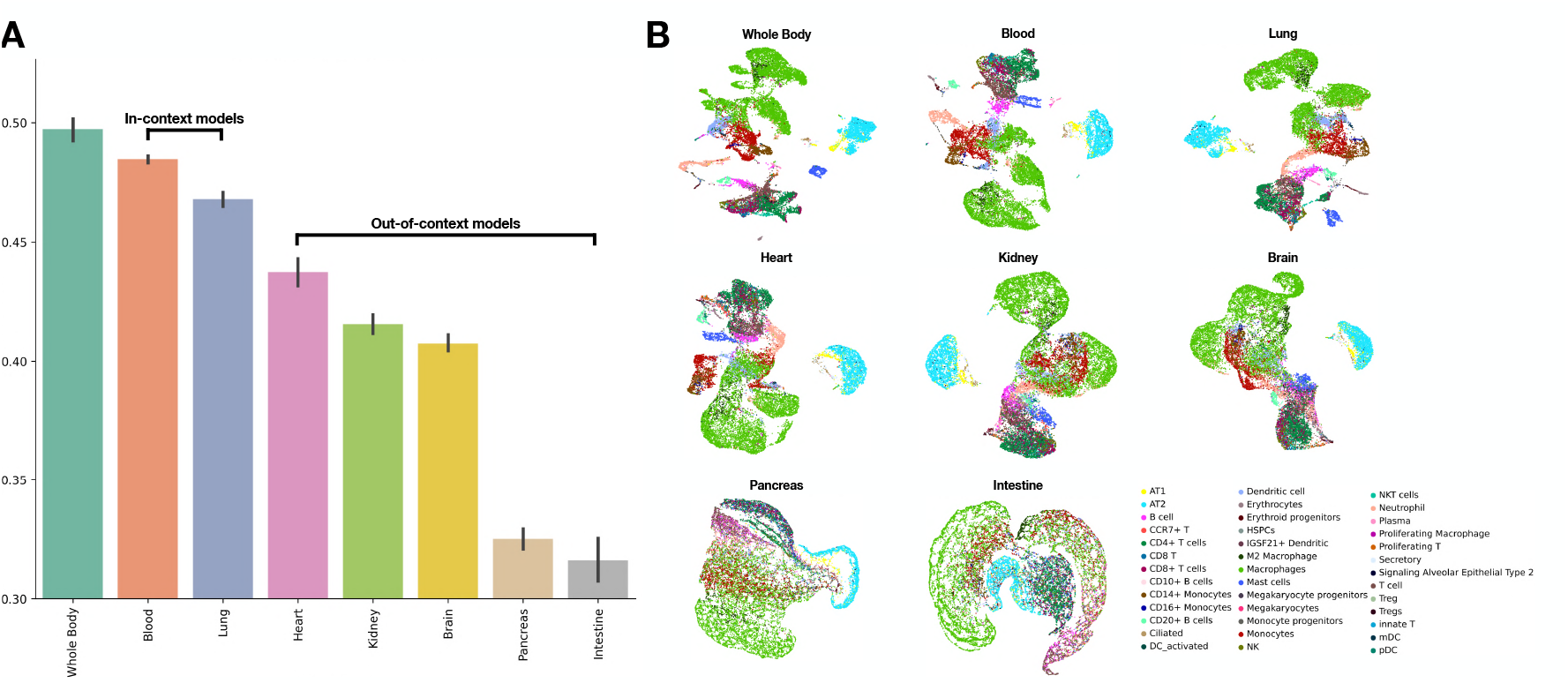
scRNA-seq batch integration results on the COVID-19 dataset (comprising 18 batches) [13], with fine-tuning applied on each of our tissue-specific pre-training models. The COVID-19 dataset comprises a range of samples, including those derived from COVID-19, lung, bone marrow, and PBMC data, reflecting its cellular context. *(A)* Illustration of the average *AvgBIO* score along with the standard error derived from five scRNA-seq integration experiments for each pre-training model. The *AvgBIO* score provides a quantitative measure of the efficacy of each pre-trained model in integrating batches. The models are ordered by their average score, highlighting the potential of tissue-specific pre-training in improving the performance of integration. In-context models (Blood and Lung), which align with the cell types in the target dataset, are separated from out-of-context models (Heart, Brain, Kidney, Pancreas, Intestine) by annotation brackets, highlighting the influence of cellular context on the integration performance. *(B)* UMAP visualization of the cell embeddings, colored by *cell types*, achieved by fine-tuning each of the tissue-specific pre-training models. The plots depict the quality of learned representations in the context of cell type diversity, offering visual confirmation of the models’ integration capabilities.

## Footnotes

1 pytorch embedding layer

2 The Census API is accessible at https://chanzuckerberg.github.io/cellxgene-census/python-api.html. It hosts and updates online data releases regularly. We used the release on May 15, 2023

